# Dynamic architecture of the Rixosome reveals mechanism of activation and ITS2 processing

**DOI:** 10.64898/2026.09.08.750072

**Authors:** Matthias Thoms, Jingyi Zhu, Xiaying Hong, Timo Denk, Jingzhi Stritzel, Shuangshuang Niu, Otto Berninghausen, Yi Li, Jiaxian Liu, Roland Beckmann, Jingdong Cheng

**Affiliations:** Gene Center and Department of Biochemistry, University of Munich LMU, Munich, Germany; Eye & ENT Hospital, Institutes of Biomedical Sciences, Fudan University, Shanghai, China; Institute of Biochemistry and Molecular Biology, Guangdong Provincial Key Laboratory of Medical Immunology and Molecular Diagnostics, the First Dongguan Affiliated Hospital, School of Basic Medical Sciences, Guangdong Medical University, Dongguan, China; Cluster for Nucleic Acid Sciences and Technologies – NUCLEAT

## Abstract

Eukaryotic ribosome assembly requires the coordinated processing and extensive remodeling of pre-rRNAs. During late nuclear maturation of the 60S subunit, sequential removal of the internal transcribed spacer 2 (ITS2) is initiated by endonucleolytic cleavage at site C_2_ by the conserved Las1 nuclease. Las1 acts together with the kinase Grc3 and the Rix1 complex to form the Rixosome, which also functions in transcriptional regulation. However, the assembly of the Rixosome, its recruitment to pre-ribosomes, and its activation for ITS2 cleavage remain unclear. Here, we present cryo-EM structures of the human LAS1 complex, two structures of the isolated Rixosome and nine transition states of Rix1-bound pre-60S particles from *Schizosaccharomyces pombe*. These structures reveal a dynamic Rixosome architecture in which the heterotetrameric Las1 complex engages one or two copies of the Rix1 complex. Rix1 binding is highly flexible in the human Rixosome but rigid in the yeast complex. The isolated yeast Rixosome remains inactive, but binding to the pre-60S particle triggers a structural rearrangement that allows for substrate engagement and activation of the nuclease. Together, our results define the dynamic architecture of the Rixosome and provide a structural framework for ITS2 processing during nuclear maturation of the eukaryotic 60S ribosomal subunit.

## Introduction

Eukaryotic ribosome biogenesis is a highly coordinated process that begins in the nucleolus, where RNA polymerase I transcribes a long polycistronic rRNA precursor (35S rRNA in yeast, 47S rRNA in human) containing the mature 18S, 5.8S and 25S/28S (yeast/human) rRNA sequences, spaced by external (5’ ETS, 3’ ETS) and internal (ITS1, ITS2) transcribed spacers^1–4^. The precursor rRNA is subsequently processed, folded, and modified, while ribosomal proteins are incorporated, and more than 200 assembly factors (AF), including RNA-binding proteins, nucleases, modification enzymes, RNA helicases and ATPases/GTPases, are dynamically exchanged to promote the maturation of the ribosomal subunits. The resulting pre-ribosomal particles undergo extensive processing and quality-control steps already in the nucleolus before maturation continues in the nucleoplasm and finally reaches completion in the cytoplasm^1–4^.

While the 5.8S and 25S/28S rRNAs are core components of the eukaryotic 60S subunit, the ITS2 (internal transcribed spacer 2) spacer region, located between the 5.8S and 25S/28S rRNA sequences, is excised during late nuclear pre-60S maturation^1–4^. The ITS2 region assembles with the factors Nsa3/Cic1/Utp30 (RL1D1 in human), Nop15 (MK67I), Rlp7/uL30 (RLP7), Nop7 (PES1) and Erb1 (BOP1) during the initial stages of pre-60S maturation in the nucleolus, forming the prominent foot-like structure, which is conserved from yeast to human ^5–13^. During the transition from nucleolar to nucleoplasmic pre-60S states, the ITS2-foot undergoes extensive remodeling that prepares it for subsequent removal. Here, the AAA-ATPase Rea1 interacts with Ytm1 through its MIDAS domain and promotes ATP-dependent release of the Ytm1-Erb1 module from the ITS2-foot structure which triggers major structural rearrangement within the pre-60S subunit^14–18^. Following this remodeling step, the Mtr4 adaptor Nop53 replaces Erb1 and becomes incorporated into the ITS2-foot region, establishing a distinct nucleoplasmic architecture that prepares the particle for subsequent ITS2 processing^19,20^.

Removal of the ITS2 is a highly coordinated process that is initiated by a conserved multienzyme machinery composed of the endonuclease Las1 and kinase Grc3 in yeast (Las1 complex), or LAS1L and NOL9 (LAS1 complex) in human ^21,22^. Las1/LAS1L is a metal-independent HEPN-family endoribonuclease containing the characteristic RφxxxH catalytic motif and dimerization juxtaposes two HEPN motifs to form the composite active site required for cleavage of ITS2 at the conserved C_2_ site in yeast (site 4 in human)^23,24^. This cleavage generates the yeast 7S and 26S precursors of the mature 5.8S and 25S rRNAs, respectively, leaving a 2′,3′-cyclic phosphate at the 3′ end of the 7S pre-rRNA and a 5′-OH at the 5′ end of the 26S pre-rRNA^21^. Grc3/NOL9 phosphorylates the newly generated 5′-OH of the 26S pre-rRNA, creating the 5′-phosphate required for 5′-3′ exonucleolytic trimming by Rat1-Rai1 (XRN2 in human) to generate the mature 5′ end of 25S/28S rRNA^21^. On the other side, processing of the 7S pre-rRNA is dependent on the nuclear exosome^25^. The N-terminal region of Nop53, which contains the conserved arch-interacting motif (AIM), is positioned close to the ITS2 region. The AIM directly binds the KOW domain of the RNA helicase Mtr4, thereby recruiting the nuclear exosome to the pre-60S particle^5,26,27^. Here, Mtr4 directs the 3′ extension of the 7S pre-rRNA toward the exosome, where the processive nuclease Rrp44/Dis3 degrades the ITS2, producing the stable 5.8S+30 intermediate. The remaining 3′ extension is subsequently handed over to the nuclear exosome-associated exonuclease Rrp6, which trims precursor further to generate the 6S pre-rRNA^20,28^, before final maturation of the 5.8S 3′ end occurs at later cytoplasmic stages^29^ (**Extended Data Fig.1**).

In both fission yeast and human cells, the Las1/LAS1 complex functions as the nuclease-kinase module of the Rixosome, which contains the Las1/LAS1 complex and the Rix1/RIX1 complex (including Rix1/PELP1, Ipi1/TEX10 and Ipi3/WDR18). The Rixosome complex further cooperates with the AAA-ATPase Rea1/MDN1, while the mammalian Rixosome additionally associates with the SUMO protease SENP3^22,30,31^. During ribosome biogenesis, the Rixosome associates with pre-60S particles and links ITS2 processing to late nuclear 60S maturation. Impairment of the ITS2 processing pathway, either through depletion or inactivation of Las1 or impaired exosome recruitment, leads to the accumulation of aberrant 80S ribosomes in the cytoplasm still harboring the ITS2-foot structure^32–34^. In addition, impairment of this pathway is linked to human diseases, as pathogenic variants in LAS1L have been associated with X-linked intellectual disability and severe congenital motor-neuron disorders with respiratory failure^35,36^. Beyond its role in ribosome biogenesis, the Rixosome is also involved in Polycomb-associated gene silencing through degradation of nascent RNAs both in fission yeast and human cells ^37,38^.

Recent structural studies have determined the molecular architecture of isolated Las1 and Rix1 complexes from multiple species, revealing the organization and catalytic regulation of these individual modules ^7,39–41^. In parallel, numerous cryo-EM structures of pre-60S particles have visualized the ITS2-foot structure, as well as later nuclear maturation intermediates bound by the Rix1 complex or the Rix1-Rea1 machinery^5–9,11–13,15,19,31,32,42–47^. However, these studies have not captured the architecture of the Rixosome in a defined functional state. In addition, how the Rixosome nuclease activity is controlled remains poorly understood. Especially, since Las1, unlike stress-responsive HEPN-family nucleases such as RNase L and Ire1^23^, forms a constitutive (Las1-Grc3)_2_ tetramer. Uncontrolled activation of such a nuclease could therefore result in inappropriate RNA cleavage, suggesting that Las1 activity must be tightly controlled until the Rixosome encounters the correct ITS2 containing pre-60S substrate^48^.

Here, we determined a series of cryo-EM structures, including the structure of the human LAS1 complex, as well as the first high-resolution structures of an isolated, complete Rixosome, and nine nuclear pre-60S states from fission yeast *Schizosaccharomyces pombe* (*S. pombe, sp)*. Our structural analysis reveals that the dimeric arrangement of the Las1 complex enables stable interaction with up to two Rix1 complexes either flexible (human) or rigid (fission yeast). The distinct conformation of the yeast Rixosome holds the complex in an inactive resting state. The orientation of the Las1 complex blocks the pre-60S interaction sites of the Rix1 submodule, and both Grc3 and Las1 remain in inactive conformations. This autoinhibitory arrangement may represent an important regulatory mechanism that ensures ribonuclease activity only to occur once the Rixosome engages with its pre-60S substrate. Further analysis of Rix1-containing pre-60S particles shows that binding of the Rixosome to its pre-60S target leads to a drastic rearrangement of the Las1 subcomplex. Together, our study reveals an unexpected multivalent organization of the Rixosome and provides a structural basis for ITS2 processing during late nuclear 60S maturation.

## Results

### Cryo-EM structure of full-length human LAS1L-NOL9 complex

The Rixosome is a conserved nuclear RNA processing machine that functions in both ribosome biogenesis and transcriptional regulation^22,37,38^, yet its overall architecture remains poorly understood. To gain structural insight into the human Rixosome, we first purified the full-length human LAS1L-NOL9 complex (**Extended Data Fig. 2a**), and tested the activity of the purified human complex using the well characterized ITS2 RNA fragment from *S. cerevisiae* as substrate^40^. The wild-type LAS1 complex efficiently cleaved the ITS2 substrate, whereas the LAS1L H160A mutant, in which the conserved catalytic histidine of the HEPN^23^ nuclease domain is mutated, failed to do so (**Extended Data Fig. 2b**). Thus, the purified sample contains an enzymatically active human LAS1 complex suitable for structural analysis.

We subjected the purified complex to cryo-EM single particle analysis and obtained a reconstruction of the human LAS1 complex at an overall resolution of 3.4 Å, which allowed us to build an improved molecular model, assisted by AlphaFold^49^ predictions (**Fig. 1a-b** and **Extended Data Fig. 2c-g, Extended Data Table 1**). Similar to previously reported structures^7,39–41^, the human LAS1 complex adopts a butterfly-like architecture containing two dimerizing LAS1L subunits and two NOL9 subunits. In detail, LAS1L can be divided into four regions: the N-terminal HEPN nuclease domain, a central helical domain (HD), the NOL9-binding motif (NBM), and the C-terminal TEX10-binding motif (TBM). In our structure, the HEPN domain and the NBM of LAS1L are well resolved, while the HD and the C-terminal TBM are not visible (**Fig. 1a-b**). However, both 2D class averages and low-pass filtering of the reconstruction revealed an additional, blurred density adjacent to the LAS1L HEPN dimer at the center of the butterfly-like particle (**Fig. 1c-d**). This density likely corresponds to the LAS1L HD, indicating that this region remains highly flexible in the human LAS1 complex. Meanwhile, NOL9 can be divided into three regions: an N-terminal domain (NTD), a central polynucleotide kinase domain (PNK), and a C-terminal domain (CTD). Two NOL9 molecules form the two wings of the butterfly-like particle, whereas the LAS1L dimer is positioned in the center (**Fig. 1a-b**).

**Figure 1.**
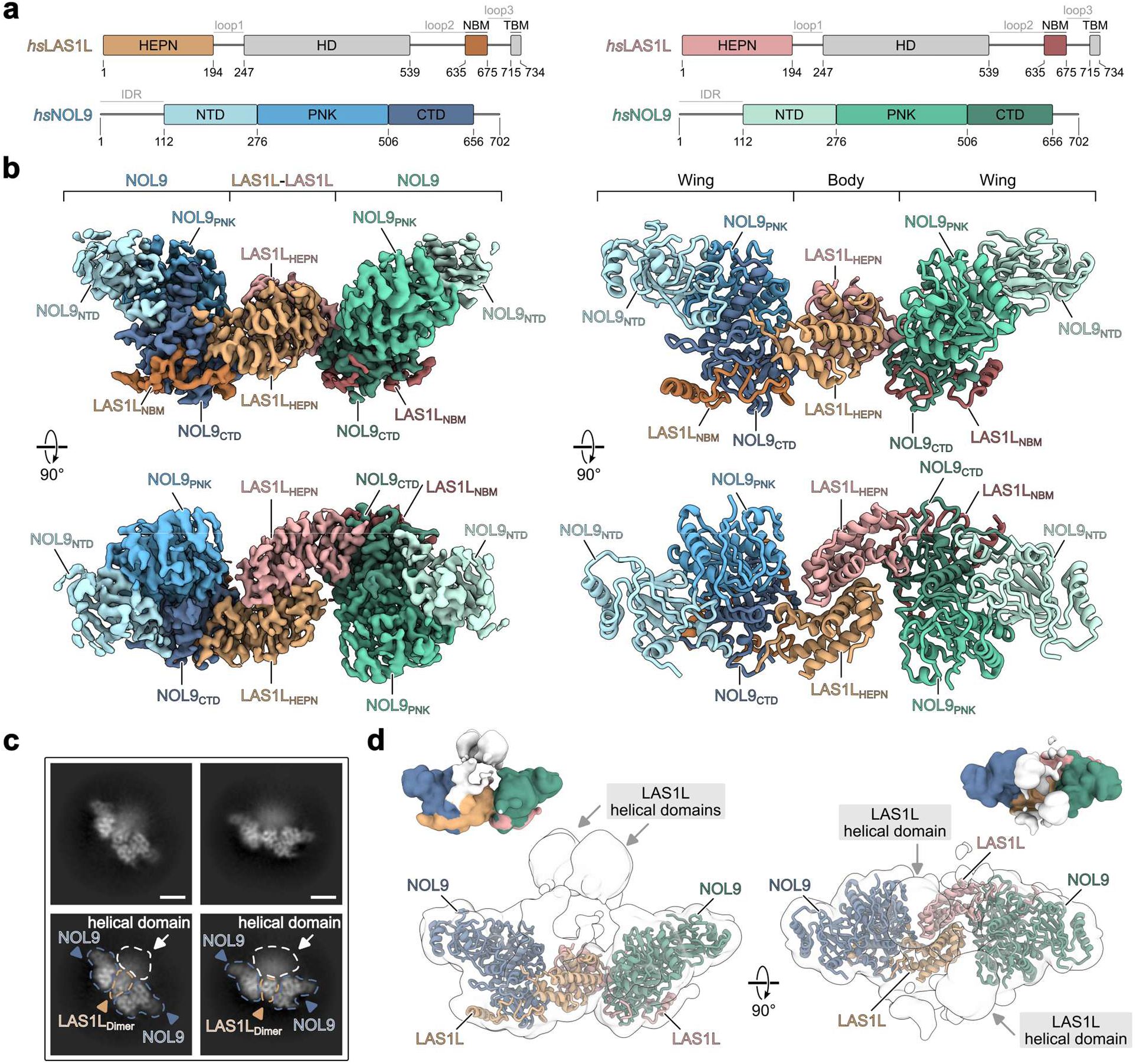
Cryo-EM structure of the full-length human LAS1 complex. **a**, Domain architecture of human LAS1L and NOL9. LAS1L contains an N-terminal HEPN nuclease domain, a helical domain (HD) and an NOL9 binding motif (NBM), as well as a C-terminal TEX10 binding motif (TBM). NOL9 contains an N-terminal domain (NTD), a polynucleotide kinase domain (PNK) and a C-terminal domain (CTD). Regions resolved in the cryo-EM structure are colored as in the model, while unresolved regions are shown in grey. **b**, Cryo-EM density map (left) and the corresponding atomic model (right) of the human LAS1 complex are shown in two orientations. **c**, Representative 2D class averages of the human LAS1 complex. The LAS1Ldimer, NOL9, and the additional central density corresponding to the flexible LAS1L helical domain are indicated. Scale bar = 50 Å. **d**, Low-pass filtered cryo-EM density showing additional central density that corresponds to the highly flexible LAS1L helical domains.

To understand how the human LAS1L and NOL9 form the nuclease-kinase submodule of the Rixosome, we focused on the well-resolved LAS1L NBM-NOL9 CTD interface (**Extended Data Fig. 3a**). In agreement with previous studies^7,39–41^, the human LAS1L NBM binds to an extended surface of NOL9. Three residues in this region, W638, W647, and L652, insert into hydrophobic pockets of the NOL9 CTD (**Extended Data Fig. 3b-c**). To test the importance of these interactions, we generated a series of LAS1L mutants and tested their ability to associate with NOL9. Single point mutations of W638 or L652 impaired LAS1L-NOL9 binding, whereas the W638A/W647A double mutant abolished the interaction (**Extended Data Fig. 3d-e**). In addition, truncation analysis showed that the LAS1L fragment containing residues 619-675 was sufficient to interact with NOL9, defining this region as the minimal NOL9-binding motif in human (**Extended Data Fig. 3d, f**).

### Dual interaction sites mediate human LAS1 and RIX1 complexes association

After solving the structure of the human LAS1 complex, we next were interested in determining how this nuclease-kinase submodule is assembled with the RIX1 complex to form the human Rixosome. To this end, we first wanted to map the interaction sites between the LAS1 and RIX1 complexes. We co-expressed Flag-ProtA-LAS1L and NOL9 with individual components of the RIX1 complex, including PELP1 (Rix1 in yeast), TEX10 (Ipi1 in yeast), and WDR18 (Ipi3 in yeast), and evaluated their association by affinity purification (**Fig. 2a**). Here, the LAS1 complex co-purified efficiently with both WDR18 and TEX10 (**Fig. 2a**, lanes 3, 4), and the complete RIX1 complex (**Fig. 2a**, lane 5), but failed to interact with PELP1 alone (**Fig. 2a**, lane 2), indicating that the human LAS1 complex contacts the RIX1 complex through WDR18 and TEX10. This result is consistent with a previous co-expression study^39^,

**Figure 2.**
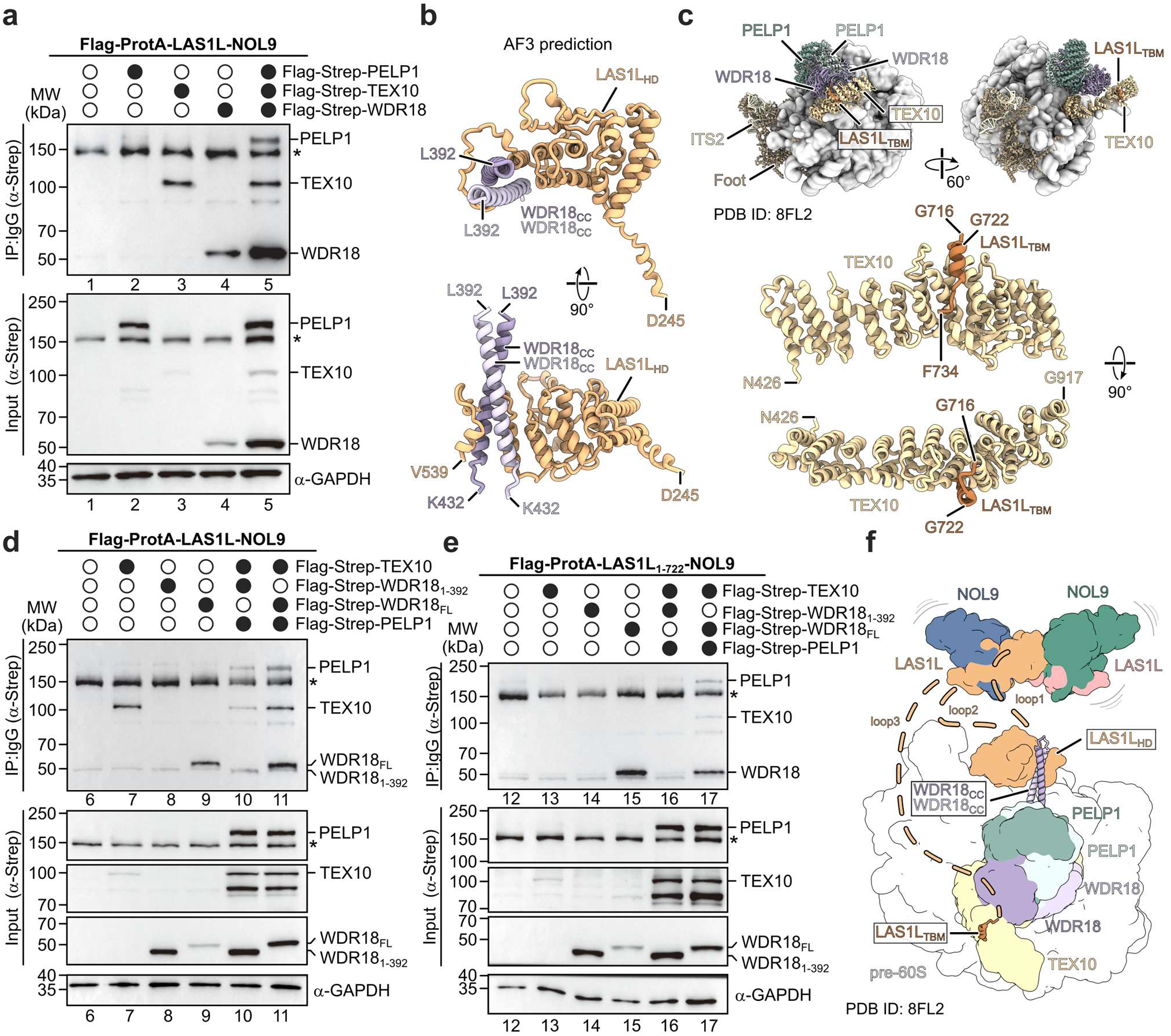
The interaction sites between human LAS1 and RIX1 complexes. **a**, Affinity purification analysis of the interaction between the human LAS1 complex and the individual components of the RIX1 complex (PELP1, TEX10 and WDR18). Flag-ProtA-LAS1L-NOL9 was co-expressed with Flag-Strep-tagged PELP1, TEX10 or WDR18, followed by IgG pulldown and western blotting. **b**, AlphaFold3 prediction of the interaction between the LAS1L HD and two copies the WDR18 C-terminal coiled coil helix (WDR_CC_, aa392-432). **c**, Structure of the human pre-60S particle bound by the RIX1 complex (PDB: 8FL2, state I1), shown in two different orientations. The RIX1 complex and foot structure are depicted as cartoon model. The lower panels show close-up views of the interaction between the LAS1L TBM and TEX10 in two different orientations. **d**, Affinity purification analysis showing that deletion of the WDR18 C-terminal helix (WDR18_1-392_) abolished the interaction between WDR18 and the LAS1 complex, whereas the mutant RIX1 complex containing the WDR18_1-392_ deletion can still associate with the LAS1 complex. **e**, Affinity purification analysis showing that simultaneous disruption of the WDR18 binding site and the TEX10 binding site within LAS1L (LAS1L_1-722_) impairs the interaction between the LAS1 and RIX1 complexes. **f**, Schematic model summarizing the binding sites between the LAS1 and RIX1 complexes. The LAS1 complex contacts the RIX1 complex through the LAS1L_HD_-WDR18_CC_ interface and the LAS1L_TBM_-TEX10 interface. Asterisks in **a, d** and **e** indicate Flag-ProtA-LAS1L, which contains a Protein A tag and is therefore detected by the antibody.

AlphaFold3 predictions of an interaction between the LAS1L HD and a complex of the two C-terminal coiled-coil helices of two WDR18 molecules, as well as with previously reported human pre-60S structures identifying the contact of the C-terminal TBM of LAS1L with a mammal-specific region of TEX10 (**Fig. 2b-c** and **Extended Data Fig. 4**)^5^.

Based on the predicted LAS1L-WDR18 interaction, we generated a WDR18 C-terminal helix deletion mutant (WDR18_1-392_), which indeed disrupts its direct interaction with the LAS1 complex (**Fig. 2d**, lane 8). However, when the WDR18 deletion mutant was present in the context of the complete RIX1 complex, the mutant complex could still associate with the LAS1 complex, albeit less efficiently than the wild-type complex (**Fig. 2d**, lane 10, 11). This indicates that the TEX10 interaction of LAS1L can compensate for the loss of the WDR18 interface in the context of the intact RIX1 complex. Therefore, we next tested the contribution of the TEX10-binding site by deleting the C-terminal TBM of LAS1L (LAS1L_1-722_, **Fig. 2e**). As anticipated, deletion of the LAS1L TBM did not affect the interaction between the mutant LAS1 complex and WDR18 (**Fig. 2e**, lane 15) and conversely, deletion of the TEX10 binding site did not abolish RIX1 complex binding (**Fig. 2e**, lane 17). Only when the LAS1L TBM deletion was combined with the WDR18 C-terminal helix deletion was the association between the LAS1 and RIX1 complexes almost completely impaired (**Fig. 2e**, lane 16). Thus, the WDR18- and TEX10-binding sites are the main interaction sites between the RIX1 and LAS1 complexes, acting independently but synergistically to mediate stable assembly of the Rixosome complex (**Fig. 2f**). However, this dual-binding mode generates a highly flexible architecture which makes structural investigation of the whole human Rixosome challenging. While the LAS1L HD and TBM engage the RIX1 complex via WDR18 and TEX10, respectively, each interaction site connects to the LAS1 complex core through long, unstructured loop regions (**Fig. 1a, 2f**). This allows the two subcomplexes to adopt variable relative orientations. Our attempts to determine the full structure of the human Rixosome were unsuccessful, likely due to this conformational flexibility.

### Cryo-EM structure of the S. pombe Rixosome

We wondered whether this highly flexible architecture of the human Rixosome is a mechanistic necessity, and therefore a feature conserved in all eukaryotes. We thus compared the protein sequences of different organisms, for a potential structural investigation of a more rigid Rixosome complex. The N-terminal HEPN domain of LAS1L is well conserved but is followed by the poorly conserved C-terminal region, including the HD and the described TBM-TEX10 interaction which is specific to vertebrates. Therefore, we focused on the HD-mediated interaction and compared the LAS1L homologs from different organisms. This analysis showed that *S. pombe* Las1 has the shortest loop regions flanking the HD (**Extended Data Fig. 5a**). AlphaFold prediction further suggested that the *S. pombe* Las1 HD is closely connected to the neighboring structured regions, making it a more compact candidate for structural analysis (**Extended Data Fig. 5b**).

Based on this analysis, we endogenously tagged the Grc3 subunit with an N-terminal TAP-Flag tag and used it as bait for affinity purification of the *S. pombe* Rixosome. The purified sample was enriched for Grc3 and Las1, as well as the components of the Rix1 complex (Rix1, Ipi3 and Ipi1) (**Fig. 3a, Extended Data Fig. 6a**), and was therefore subjected to cryo-EM single-particle analysis (**Extended Data Fig. 6**). 2D classification revealed particles with a two-lobed architecture, in which one or two ball-like Rix1 complexes are connected to the central Las1 complex consisting of two copies of Las1 and Grc3 (**Fig. 3b-c, Extended Data Fig. 6c**). In addition to these Rixosome particles, a small population of pre-60S particles was also present (see below, **Extended Data Fig. 6d-e**). After heterogeneous refinement, 3D classification and non-uniform refinement of the Rixosome particles, we obtained two major states of the complete Rixosome at average resolutions of 2.7 Å and 2.9 Å, respectively. The first state (single Rix1 Rixosome) contained a single Rix1 complex rigidly associated with the central Las1 complex, while the second state (double Rix1 Rixosome) contained two clearly resolvable Rix1 complexes bound to the Las1 complex on opposite sides. This finding indicates that the Rixosome can act in two different modes, with either one or two Rix1 complexes integrated, and that the high degree of flexibility as observed in the human system is indeed not a conserved feature. Local refinement of the individual Rix1 and Las1 complexes further improved the quality of the maps and allowed us to build molecular models of both *S. pombe* Rixosome assemblies (**Extended Data Figs. 7-8, Extended Data Table 2, Extended Data Movie 1**).

**Figure 3.**
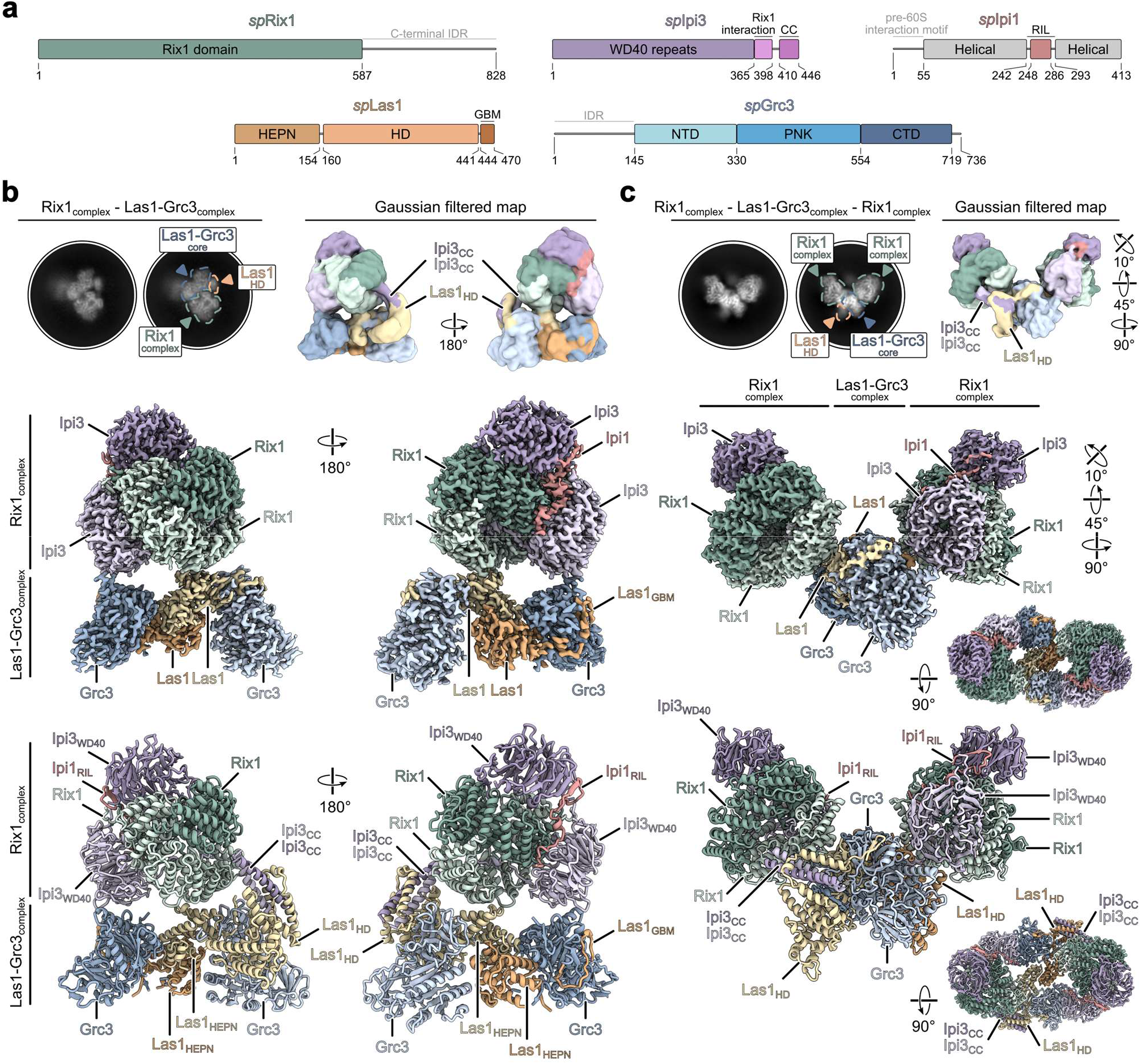
Cryo-EM structures of the S. pombe Rixosome. **a**, Domain architecture of *S. pombe* Rix1, Ipi3, Ipi1, Las1 and Grc3. Rix1 contains an N-terminal Rix1 domain and a C-terminal intrinsically disordered region (IDR). Ipi3 contains WD40 repeats and a C-terminal coiled-coil (CC) region. Ipi1 contains an unstructured N-terminal pre-60S binding motif and a Rix1-interacting-loop (RIL) inserted into the helical domain. Compared with the human LAS1L, *S. pombe* Las1 lacks the C-terminal TEX10 binding motif (TBM). **b-c**, Cryo-EM structures of the single Rix1 Rixosome (**b**) and the double Rix1 Rixosome (**c**). Representative 2D class averages and Gaussian-filtered maps are shown at the top. The Gaussian filtered maps show the density of the Las1 HD bound to two Ipi3 coiled-coil helices. Cryo-EM density maps (middle) and atomic models (bottom) are shown in two different views for each structure.

Each Rix1 complex consists of two Rix1, two Ipi3 and one Ipi1 subunit, forming a globular, ball-like structure as was previously observed for Rix1 complexes from *S. cerevisiae, C. thermophilum*, and humans^5,7,19,31,39,50^. In our reconstruction, only the Rix1 domain of Rix1, nearly full-length Ipi3, and the Rix1-interacting loop (RIL) of Ipi1 could be clearly resolved. Due to the flexibility, the N-terminus and helical domain of Ipi1, both of which are required for the stable binding of the Rix1 complex to pre-60S subunits, were not visible. Nevertheless, at a lower contour level, we observed a poorly resolved globular density between the two Rix1 complexes that likely corresponds to the helical domains of the Ipi1 subunits (**Extended Data Fig. 6e**, right panel *). Similar to the human LAS1 complex, the *S. pombe* Las1 complex also consists of two Las1-Grc3 heterodimers that form the conserved butterfly-like architecture by dimerizing their Las1 HEPN domains. In the single Rix1 Rixosome assembly, one Rix1 complex binds to one of the two Las1-Grc3 heterodimers, resulting in an asymmetric Rixosome structure. In contrast, within the double Rix1 Rixosome complex, two Rix1 complexes bind to opposite sides of the bridging Las1 complex. Each Rix1 complex engages one Las1-Grc3 heterodimer, forming a pseudo-symmetrical, twisted, C-shaped assembly. Thus, the *S. pombe* Rixosome can exist in both one and two Rix1 complex forms. For both Rixosome states we observe the flexible but defined density of the Las1 HD-Ipi3 CC interaction, consistent with AlphaFold predictions and previous observations^7,39^ (**Fig. 3b, c**, upper panels; **Extended Data Fig. 7 and 8a, Extended Data Movie 1**).

We next compared the human LAS1L-NOL9 complex with the *S. pombe* Las1-Grc3 complex and previously reported fungal Las1-Grc3 structures (**Extended Data Fig. 9**)^7,39–41^. All complexes adopt a conserved butterfly-like architecture, indicating that the overall organization of the LAS1/Las1 nuclease-kinase module is conserved from fungi to human. However, the human LAS1 complex displays a more open arrangement between the two wings formed by the two human NOL9 subunits, when compared with the fungal complexes, including the *S. pombe* structure determined here (**Extended Data Fig. 9**). This conformational difference may reflect species-specific adaptations of the LAS1 complex for processing ITS2 regions of different lengths and shapes.

### The S. pombe Las1 HD defines the major assembly interface of the Rixosome

We next analyzed how the Las1 and Rix1 complexes are associated within the assembled *S. pombe* Rixosome. In agreement with our human interaction mapping (**Fig. 2**), the major interface between the Las1 and the Rix1 complex is mediated by the Las1 HD and the C-terminal helices of two Ipi3 molecules, together with two additional unforeseen contact sites (**Fig. 4, I-IV**, **Extended Data Fig. 8a)**.

The Las1 HD is positioned on the outer side of the central Las1 complex, directing interaction with one of the Grc3 copies and protruding toward the neighboring Rix1 complex (**Fig. 4**). Here, it directly forms an extended contact surface with a composite Ipi3 C-terminal coiled-coil structure, which is formed by the intertwined C-terminal helices from the two Ipi3 subunits within one Rix1 complex (**Fig. 4**, III-IV). This arrangement is consistent with AlphaFold prediction and corresponds to the conserved LAS1L HD-WDR18 interaction identified in the human system^39^ (**Fig. 2, Extended Data Figs. 4b** and **8a**). Besides this major interface, two additional contacts were observed between the Las1 and Rix1 complexes from *S. pombe*: I) Grc3 interacts with one of the Ipi3 WD40 domains and II) the Las1 HEPN region contacts one of the Rix1 molecules within the Rix1 complex (**Fig. 4**, I-II). These auxiliary interfaces may further fine-tune the relative orientation between the Las1 and Rix1 complexes (see below).

**Figure 4.**
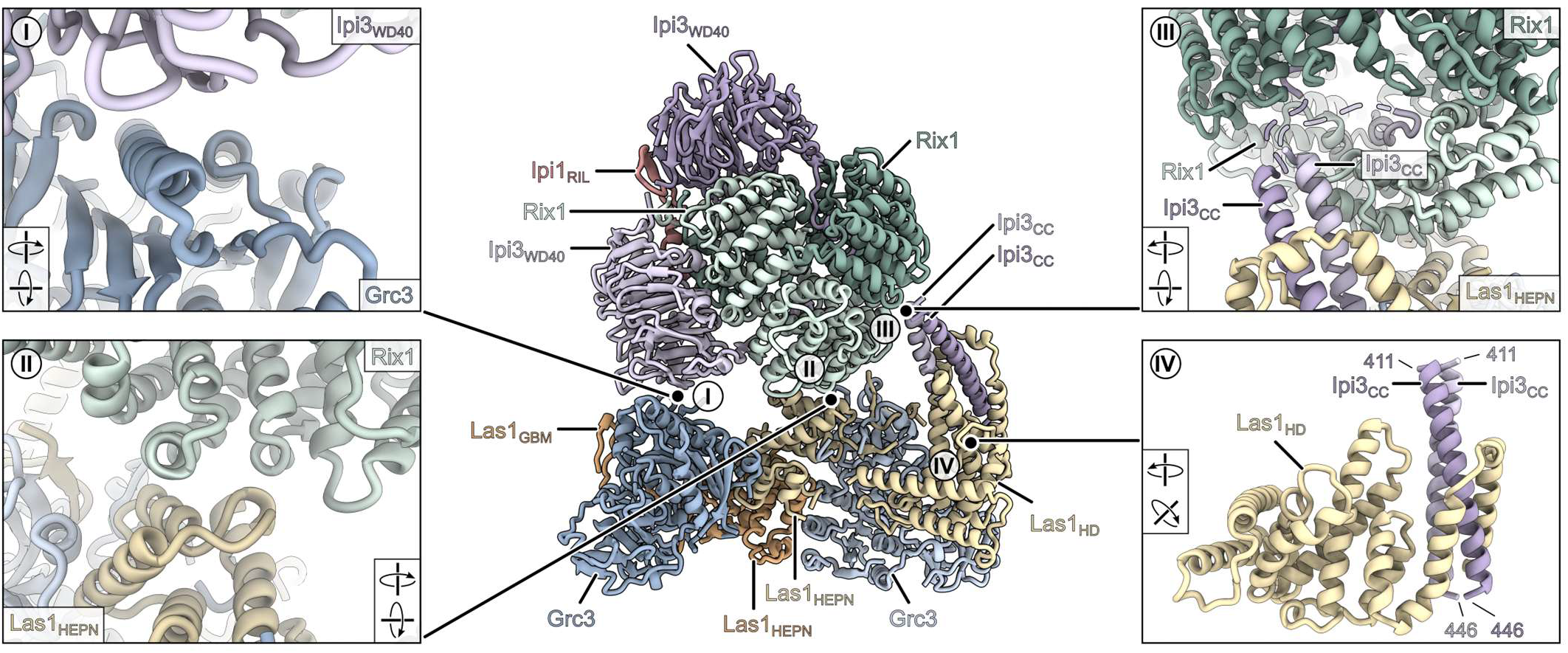
Interfaces between the Las1 and Rix1 complexes. Close up views of the interfaces between the Las1 and the Rix1 complexes in the *S. pombe* Rixosome. The central panel shows the overall structure of the single Rix1 Rixosome, with boxed regions indicating the contact sites. Panels **I** and **II** show auxiliary interfaces between Grc3 and the Ipi3 WD40 domain (**I**) and between Las1 HEPN and Rix1 (**II**). Panels **III** and **IV** highlight the major interface between the Las1 HD and the Ipi3 CC region. The missing connection between the Ipi3 CC region and the Ipi3 WD40 domain within the Rix1 complex is shown as dashed lines.

To understand the catalytic state of the Rixosome, we then analyzed the nuclease and kinase centers within the Las1 complex. In the Las1 nuclease center, the conserved HEPN catalytic motif is well resolved. Here, the histidine switch residues are not arranged towards the nuclease active site as reported for the active RNase state^41^. Instead, they adopt a conformation similar to the previously described inactive resting state (**Extended Data Fig. 10a**)^41^. Consistently, no nucleotide density was observed in the Grc3 active site. In addition, the P-loop and lid regions are positioned in a closed conformation, rather than the open conformation required for ATP binding and catalysis (**Extended Data Fig. 10b**). Thus, both the Las1 nuclease center and the Grc3 kinase center in the free *S. pombe* Rixosome are in an catalytic inactive state. This resting conformation may prevent nonspecific RNA cleavage before the complex engages the pre-60S particle and is in agreement with studies in budding yeast showing the direct coupling of nuclease and kinase activity^51,52^.

### Cryo-EM structures of Rix1-associated S. pombe pre-60S ribosomal subunits

We next aimed to explore how the Rixosome functions on pre-60S particles during ITS2 processing. For this purpose, we tagged the Rix1 subunit of the Rix1 complex with a C-terminal FTpA tag in *S. pombe* and used it as bait to purify endogenous nuclear pre-60S particles and subjected the purified sample to cryo-EM single particle analysis. 2D classification identified mainly pre-60S particle classes but also free Rixosome and Rix1 complex class averages (**Extended Data Fig. 11a-c**). The pre-60S classes were highly heterogeneous, and extensive 3D classification resolved nine distinct pre-60S states, with an overall resolution ranging from 2.4 Å to 2.9 Å (**Fig. 5, Extended Data Figs. 11-15, Extended Data Table 3**). Based on the maturation status of the rRNA and bound AFs, these particles could be broadly classified into three major groups, referred to as states A, B and C (**Fig.5, Extended Data Fig. 15a**).

**Figure 5.**
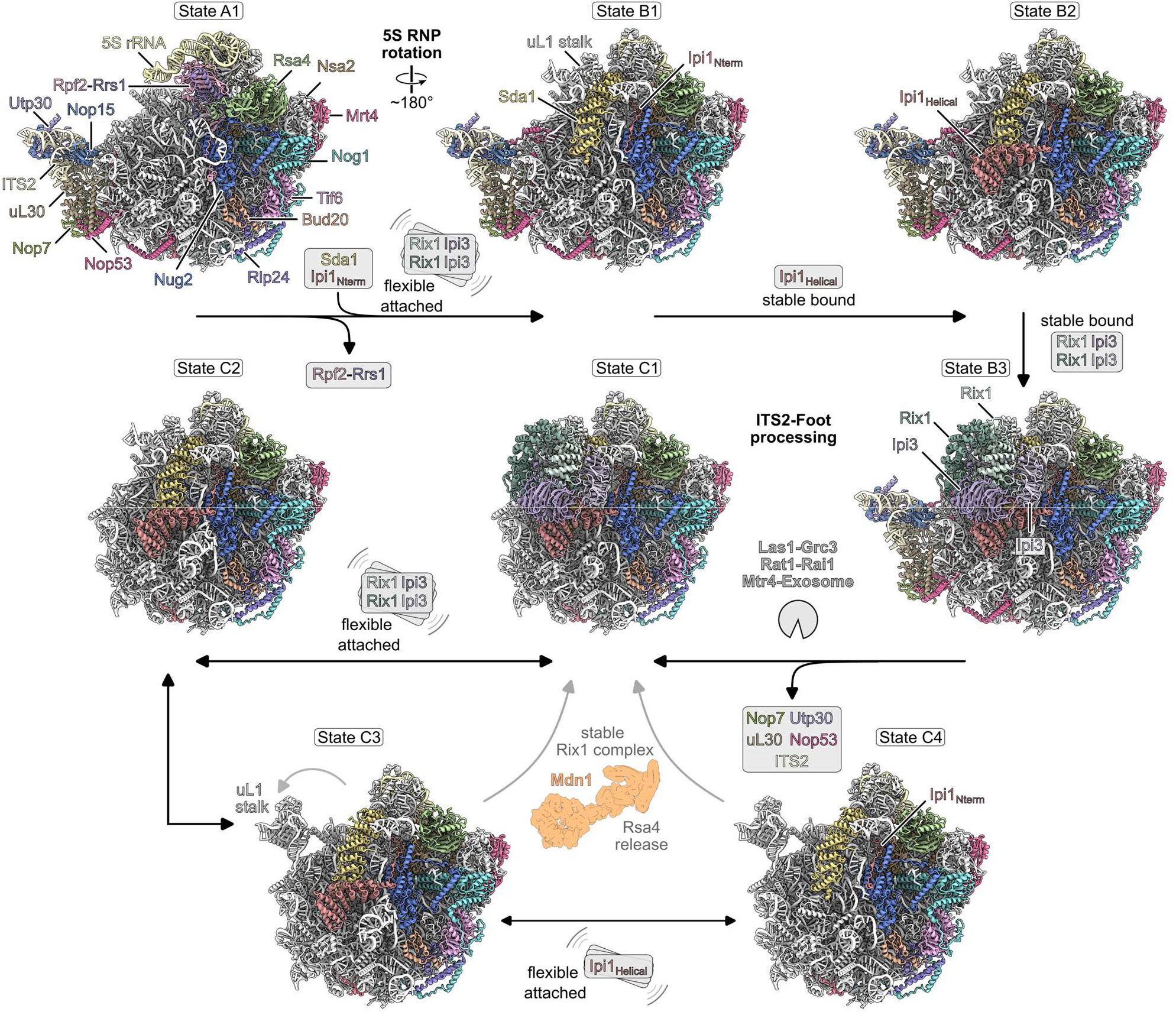
Cryo-EM structures of Rix1 bound pre-60S ribosomes. Cryo-EM structures of *S. pombe* pre-60S particles purified using Rix1 as bait. The particles are classified into three major groups (states A, B and C) based on their maturation features. State A represents an early nucleoplasmic pre-60S particle with an intact ITS2-foot structure and the 5S RNP in the pre-rotation stage. The state B substates represent pre-60S particles with progressive stabilization of the Rix1 complex (states B1-B3). The different C substates represent later pre-60S particles after ITS2-foot processing with differences in the interaction of the Rix1 complex with the pre-60S moiety (C1-C4). Black arrows indicate the proposed maturation trajectory during ITS2 processing. Double headed arrows indicate that the corresponding states may interconvert, whereas grey arrows indicate possible alternative maturation paths. As indicated, the successive progression of the pathway requires the stable association of the Rix1 complex and its interaction partner, Mdn1, for the release of Rsa4.

State A represents an early group of Rix1-associated pre-60S particles with an intact ITS2-foot structure. In these substates, the foot-associated factors Nop7, Nop53, Utp30 (Nsa3/Cic1 in budding yeast) and uL30 (Rlp7 in budding yeast) are clearly visible together with the ITS2 pre-rRNA. The 5S RNP remains in an immature pre-rotation conformation, and the AFs Rpf2-Rrs1 are associated and chaperone the 5S RNP (**Fig. 5**, upper left). These features are consistent with previously described nucleoplasmic pre-60S intermediates including state B’ in human cells (PDB ID: 8IR3), the Nog2 particle in *S. cerevisiae* (e.g. PDB ID: 3JCT) as well as the state 1 in *C. thermophilum* (PDB ID: 8PV7) (**Extended Data Fig. 16**)^7,11,43^. State A can be further divided into subclasses according to the presence or absence of the ES27 interacting with Arx1 (State A1: combined state A particle, whereas state A2 contains Arx1 and bound ES27) (**Extended Data Figs. 11d** and **15b**).

State B represents a group of substates (B1-B3) in which the AFs Rpf2-Rrs1 are released and the 5S RNP has rotated into an almost mature conformation (**Fig. 5**). Within the different substates, the Rix1 complex becomes progressively stabilized on the pre-60S particle. In state B1, Sda1, the Rix1 complex interacting AF, is already positioned between the uL1 stalk and the CP density, while the Rix1 complex is visible but remains highly flexible. Only the N-terminus of Ipi1, which is required for recruitment of the Rix1 complex to pre-60S particles, can be clearly modeled^19^. In the subsequent state B2, the helical domain of Ipi1 becomes stably incorporated and interacts with Sda1. Finally, in state B3, the globular (Rix1-Ipi3)_2_ structure of the Rix1 complex is stably positioned, closely resembling previously reported states in budding yeast (Rix1-Rea1 particle, PDB ID: 6YLH), human cells (state I1, PDB ID: 8FL2), and *C. thermophilum* (state 3, PDB ID: 8PV6) (**Extended Data Fig. 16**) ^*5,7,19*^. Thus, the B states capture a series of intermediates in which Rix1 complex binding becomes stabilized and the Rixosome-associated ITS2-processing machinery is positioned on the pre-60S particles.

The subsequent state C represents a series of substates (C1-C4), in which the ITS2-foot region has already been processed (see **Fig. 5, Extended data Fig. 15a**). Compared to states A and B, the densities for ITS2 and foot-associated factors are absent, indicating that the Rixosome and the nuclear exosome have processed the ITS2. The C states can be divided into four subclasses. State C1 directly follows state B3 and represents the first ITS2-processed state. In this state, the ITS2-foot structure is removed, but the remaining pre-60S architecture remains largely unchanged, including a stable bound Rix1 complex. This is consistent with the previously reported states, including the foot removed state I3 in human (PDB ID: 8FL4), the Rix1-Rea1 particle in *S. cerevisiae* (PDB ID: 6YLH), and state 4 in *C. thermophilum* (PDB ID: 8PV8) (**Extended Data Fig. 16**)^5,7,19,45^. In the following state C2, the globular (Rix1-Ipi3)_2_ structure of the Rix1 complex becomes flexible, but remains associated with the pre-60S particle through Ipi1. State C3 differs from state C2 mainly by a rearrangement of the L1 stalk into an out position. Sda1 remains embedded in the pre-60S particle, interacting with the CP and the Ipi1 helical domain. In the final state C4, Sda1 and the Ipi1 N-terminus remain stably bound to the pre-60S particle, while the helical Ipi1 domain becomes flexible (**Fig. 5, Extended data Fig. 15a**).

This apparent reversal from a stable Rixosome-bound state in C1 to more flexible Rix1-associated states in C2-C4 may reflect the beginning of Rixosome disengagement after ITS2 cleavage and foot removal. In this scenario, the Rixosome does not immediately dissociate from the particle, but remains flexibly associated before maturation continues. As long as the Ipi1 N-terminus and Sda1 are bound to pre-60S particles, stable reassociation of the Rix1 complex is possible, which at least in budding yeast is required for the interaction with Rea1/MDN1 during the later Rsa4/NLE1 release step ^19,31,53^. However, we cannot exclude that the purification procedure may have contributed in part to this flexible association or that alternative parallel maturation routes exist. Nevertheless, the overall order of the major A, B and C groups likely reflects the main direction of pre-60S maturation captured in our *S. pombe* dataset, consistent with the conserved maturation trajectory of late pre-60S maturation described for other species^1–3^. Substages similar to the here described states B1 and C4 were recently also observed in the human system^11^.

### Pre-60S engagement induces Rixosome rearrangement for ITS2 processing

Among the Rix1-associated pre-60S intermediates, the stable Rix1-bound state B3 enabled us to study the positioning of the Rixosome on pre-60S particles immediately prior to ITS2 processing. In this state, as shown for other species, the Rix1 complex contacts the pre-60S ribosome through distinct interfaces, including interactions with the AF Sda1, the uL1 stalk and the immature helix 38 (H38)^5,7,19,31^. This binding mode places the *S. pombe* Rix1 complex at the inter subunit surface of the pre-60S subunit, adjacent to the ITS2-foot region, where the Las1 complex is expected to function.

Next, we assessed if the structure of the free Rixosome could be directly fitted onto the Rix1 complex when bound to the pre-60S ribosome in state B3. For this purpose, we superimposed the single Rix1 Rixosome (free Rixosome structure) onto the pre-60S structure by aligning the Rix1 complex and the Ipi1 RIL within the Rix1 complex. The fitting placed the Las1 complex against the pre-60S body and generated extensive steric clashes with the pre-60S particle (**Fig. 6a**). Detailed inspection showed that, in state B3, the corresponding surfaces of the Rix1 complex are occupied by 25S rRNA H38 and the uL1 stalk. In contrast, in the free Rixosome structure, the same region is contacted by Grc3 and the HD and HEPN domains of Las1 (**Fig. 6b**). Thus, the observed free Rixosome structure is incompatible with direct pre-60S binding. This indicates that the Las1-Grc3 module must undergo extensive rearrangements when transitioning from the free, resting Rixosome state to an active state associated with the pre-60S substrate.

**Figure 6.**
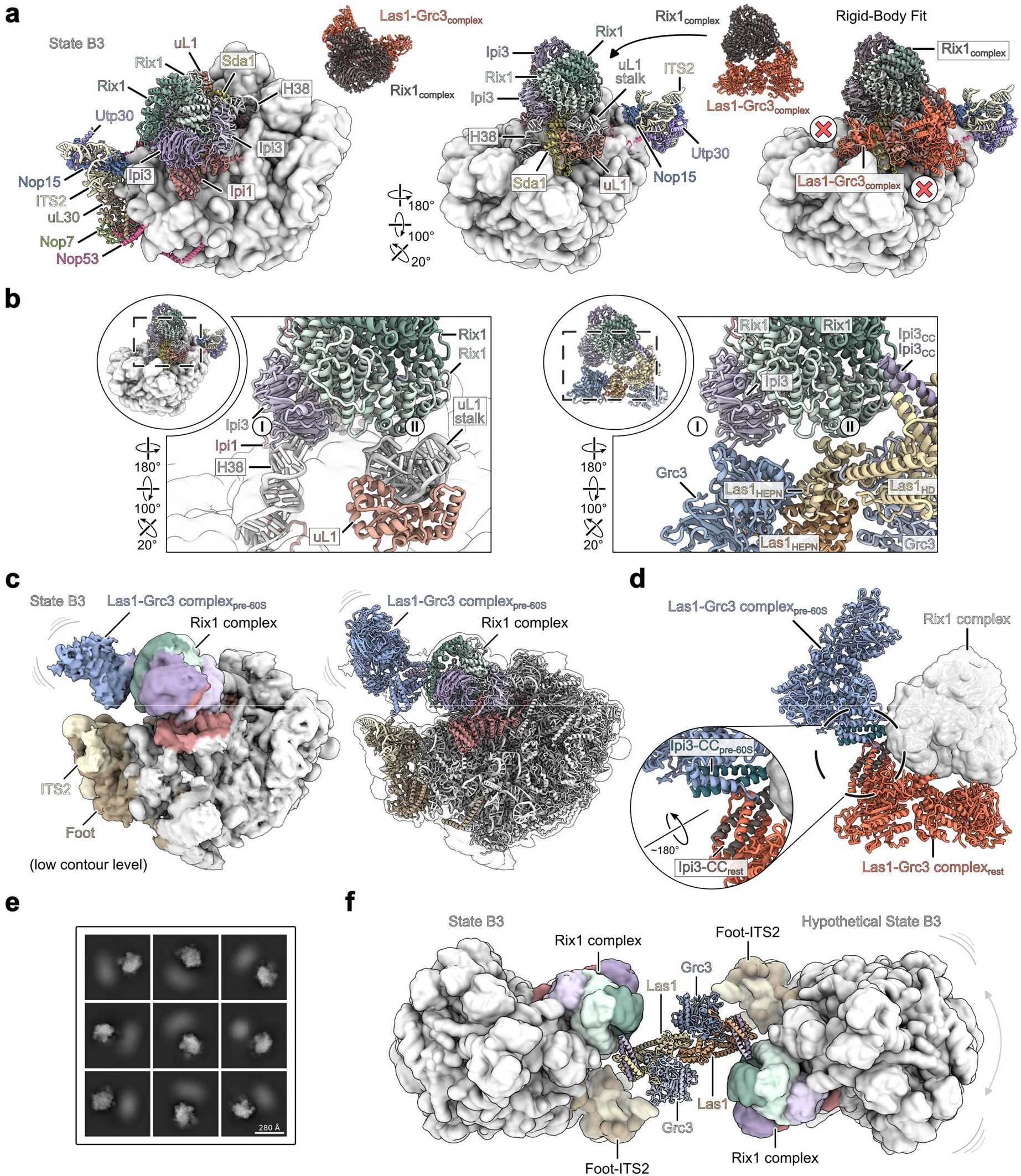
Pre-60S engagement remodels the Rixosome for ITS2 processing. **a**, Structure of the pre-60S state B3 highlighting the position of the Rix1 complex in two different orientations (left and middle). Rigid body fitting of the single Rix1 Rixosome structure onto the pre-60S state B3 results in steric clashes of the Las1-Grc3 complex with the pre-60S subunit, indicated by red cross marks (right). **b**, Close-up views and comparison showing that the Rix1 complex surface interacted with the Las1-Grc3 complex in the free Rixosome (right) is occupied by H38 and the uL1 stalk of the pre-60S ribosome in state B3 (left). **c**, Cryo-EM density map displayed at low contour level (left) and structural model (right) showing the Las1-Grc3 complex fit into the additional density in state B3. The Las1-Grc3 density is connected to the Rix1 complex and in close proximity to the ITS2-foot. **d**, Comparison between the free rest state (red) and the pre-60S bound state (blue) of the Las1-Grc3 complex based on the Rix1 complex. The Las1 complex is rotated by about 180° around the Ipi3 CC region. **e**, Representative 2D class averages of paired pre-60S particles showing additional density adjacent to the main pre-60S ribosome. **f**, Proposed model of Rixosome mediated pre-60S dimerization. One Las1 complex bridges two Rix1-bound pre-60S ribosomes and positions the two ITS2-foot regions near the Las1 catalytic centers.

Consistent with this idea, when the state B3 map was displayed at a low contour level, an additional low-resolution density was observed in direct contact with the Rix1 complex and close to the ITS2-foot region (**Fig. 6c**). Although this density was weak, its size and shape can accommodate the Las1-Grc3 complex. In this fitted position (termed pre-60S conformation), the Las1-Grc3 complex is placed right next to the ITS2-foot, with the Las1 nuclease module oriented toward the ITS2 C2 cleavage site. Comparison between the free resting state and the pre-60S bound conformation shows that this rearrangement can be achieved only by an approximately 180° rotation of the Las1 complex around the Ipi3 C-terminal coiled-coil region (**Fig. 6d**). This movement preserves the conserved Las1 HD-Ipi3 coiled-coil interaction, while relocating the Las1-Grc3 catalytic module away from the pre-60S surface and toward the ITS2-foot region. The pre-60S bound conformation may also provide a structural basis for pre-60S dimerization. In this conformation, one Las1-Grc3 complex could remain connected to the first Rix1 complex, whereas the opposite side of the dimeric Las1 complex would be exposed and could accommodate a second Rix1 complex. This second Rix1 complex would be positioned on the opposite side of the Las1 complex and could bind to another pre-60S particle in a similar manner. In such an arrangement, the dimeric Las1-Grc3 complex would bridge two Rix1-bound pre-60S particles, placing the ITS2-foot regions from both particles close to the two catalytic centers of the Las1 dimer. To investigate the presents of such a higher-order assembly in our sample, neighboring pre-60S particles within a maximum distance of 350 Å were paired based on spatial proximity. 2D class averages of paired particles frequently showed an additional density adjacent to the main pre-60S particle, likely consistent with the presence of a second but flexibly associated pre-60S particle (**Fig. 6e-f**). Together, these observations suggest that the double Rix1 Rixosome may potentially induce pre-60S dimerization in cells, thereby enabling the dimeric Las1-Grc3 complex to eventually process ITS2 more efficiently.

## Discussion

In this study, we show that the Las1 complex forms a conserved dimeric nuclease module from fungi to human. In *S. pombe*, the dimeric Las1-Grc3 complex associates with up to two Rix1 complexes to form a twisted C-shaped Rixosome, yet adopting an inactive resting conformation. Analysis of Rix1 complex-bound pre-60S structures reveals that this inactive state is relieved upon pre-60S engagement, pointing to a substrate-coupled mechanism for regulating ITS2 processing.

This substrate-coupled mechanism distinguishes Las1 from other HEPN-family ribonucleases. In general, HEPN ribonucleases require dimerization to fold and position their composite active sites, and only become catalytically active upon dimer formation. Well-characterized examples such as Ire1 and RNase L dimerize in direct response to unfolded proteins and second messengers, respectively^23^. Las1 differs, as its HEPN nuclease module is constitutively dimeric, and therefore cannot rely on nuclease dimerization itself as the principal regulatory switch. Our structural comparison of the free and pre-60S-bound Rixosome indicates that the free Las1-Grc3 module nevertheless remains catalytically inactive, and that engagement with the pre-60S particle itself might provide the activating cue that drives the Las1-Grc3 module into a conformation competent for ITS2 cleavage. Meanwhile, our *S. pombe* Rixosome structure shows that the two sides of the Las1-Grc3 dimer can each bind to one Rix1 complex, thereby creating a platform capable of engaging two pre-60S ribosomes simultaneously. This raises the possibility that the Rixosome could, in principle, coordinate two pre-60S particles during ITS2 processing, although the functional significance of this arrangement remains currently unclear. During the preparation of this manuscript, a recent in-cell cryo-ET study of human nucleoli showed that human pre-60S ribosomes also dimerize, bridged by a shared LAS1L-NOL9 heterotetramer^54^. This observation is in agreement with our *S. pombe* model and suggests that Rixosome-mediated pre-60S dimerization may be conserved from fungi to human.

In our *S. pombe* pre-60S dataset, the state A to C order likely reflects the primary direction of ITS2-foot maturation. However, this overall order does not require every subclass to follow a strictly linear pathway. Previous structural and functional studies suggest that ITS2 processing can be partially uncoupled from other late maturation events. For example, 5S RNP rotation and ITS2-foot removal are not strictly co-dependent, and pre-60S particles can proceed to later maturation stages when ITS2 processing is impaired, as in the case of Las1 depletion or impaired Mtr4-nuclear exosome recruitment^32–34^. Thus, while our structures represent the principal ITS2-processing trajectory, alternative or branched maturation routes may coexist as well.

The flexibly associated Rix1 state C may reflect this complexity. One possibility is that the flexibility of the Rix1 complex after ITS2-foot removal is partially caused by purification. Alternatively, this dynamic association may represent a genuine post processing state in which the Rixosome is still loosely bound and does not immediately dissociate from the pre-60S particle. This interpretation fits with the broader idea that the Rixosome (including Rea1/MDN1) engage pre-60S particles multiple times during maturation. The AAA-ATPase Rea1/MDN1 is known to participate, together with the Rix1 complex, in both early Ytm1 removal and later Rsa4 release^14–18,53^. Although the structural basis for the early Ytm1-removal event has not yet been directly visualized, these observations point towards the possibility that also the complete Rixosome could act repeatedly throughout 60S biogenesis: first during Ytm1 removal, later during ITS2-foot processing, and Rsa4 release.

Together, our study supports a model in which the Rixosome restricts ITS2 cleavage to correctly assembled pre-60S particles through a substrate-coupled activation mechanism, thus providing a structural checkpoint for ITS2 processing and linking ITS2 removal to the dynamic remodeling pathway of late pre-60S maturation.

## Methods

### Human Expi293F cell culture

Human Expi293F suspension cells (A14527, Thermo Fisher Scientific Inc.) were cultured in SMM 293-TII Expression Medium (M293TII, Sino Biological Inc.) under serum-free suspension culture conditions. The cells were routinely grown in 250ml glass bottles, and the medium volume was kept equal to or less than 1/3 of the total bottle volume. The bottles were fixed to a shaker (MQD-A1C, Union Bio) supplied with 5% CO_2_, and shaken at 120 rpm. Cells were maintained at a density ranging from 1 × 10^6^ cells ml^−1^ to 3 × 10^6^ cells ml^−1^. For passaging, cells were collected by centrifugation and resuspended at the desired density in fresh medium.

### Plasmid construction

The full-length coding sequences of human LAS1L, NOL9, TEX10, WDR18 and PELP1 were amplified from a human cDNA library. For cloning, a 15 bp vector homologous sequence and a KpnI site (5′ GGTACC 3′) were added to each forward primer, and a 15 bp vector homologous sequence and an XhoI site (5′ CTCGAG 3′) were added to each reverse primer. PCR was performed using PrimeSTAR Max DNA Polymerase (R045, Takara) according to the manufacturer’s instructions. The PCR products were inserted into KpnI and XhoI linearized vectors by homologous recombination. All plasmids were verified by DNA sequencing.

### Protein binding assay

For LAS1 complex and RIX1 complex binding assay, full length LAS1L and LAS1L (1-722) were cloned into the pMLink vector as previously described^55^. They were expressed with an N-terminal FLAG-4xProtein A-SUMO tag followed by TEV and 3C protease cleavage sites. NOL9 was cloned into a pMLink vector without any tag. Full length PELP1, TEX10, WDR18 and WDR18 (1-392) were cloned into the pcDNA5 FRT/TO vector with an N-terminal FLAG-Strep-II tag followed by an HRV 3C protease cleavage site. For each transfection, 12 μg of each plasmid was used, and the total amount of plasmid DNA was adjusted to 60 μg with empty pcDNA5 FRT/TO vector.

For LAS1L mutants and NOL9 binding assay, full length LAS1L, LAS1L truncations (1-619, 1-722) and LAS1L point mutants (W638R-W647R, W638A-W647A, W638R, W638A, L652R, L652A) were cloned into pMLink vector with an N-terminal FLAG-4xProtein A-SUMO tag followed by TEV and 3C protease cleavage sites. NOL9 were cloned into pcDNA5 FRT/TO vector with an N-terminal FLAG-Strep II tag and an HRV 3C protease cleavage site. For each transfection, 30 μg of each plasmid was used, and the total amount of plasmid DNA was adjusted to 60 μg with empty pcDNA5 FRT/TO vector.

For NOL9 and LAS1L truncations binding assay, NOL9 was cloned into pMLink vector with an N-terminal FLAG-4xProtein A-SUMO tag followed by TEV and 3C protease cleavage sites. Full length LAS1L and LAS1L fragments (619-675, 619-734, 636-675) were cloned into pcDNA5 FRT/TO vector with an N-terminal FLAG-Strep II tag and an HRV 3C protease cleavage site. Full length LAS1L was also cloned into the pcDNA5 FRT/TO vector with a C-terminal FLAG-Strep II tag followed by an HRV 3C protease cleavage site. For each transfection, 30 μg of each plasmid was used, and the total amount of plasmid DNA was adjusted to 60 μg with empty pcDNA5 FRT/TO vector.

The indicated plasmids were transfected into 50 ml Expi293F cell suspension at a density of 2 × 10^6^ cells ml^−1^ using PEI (40,000 MW, Yeasen). For protein expression, cells were cultured in SMM 293-TII medium supplemented with streptomycin and penicillin for 48 h. Cells were harvested by centrifugation at 3,000 rpm for 10 min and resuspended in lysis buffer containing 30 mM HEPES (pH 7.4), 150 mM NaCl, 10% glycerol, 0.2 mM EDTA, 0.25% CHAPS, 5 mM MgCl_2_, 5 mM ATP, 2 mM DTT, and protease inhibitors (1 μg ml^−1^ aprotinin, 1 μg ml^−1^ pepstatin A, 1 μg ml^−1^ leupeptin, and 0.1 mM PMSF). Cells were lysed at 4 °C for 30 min with gentle rotation. Cell debris was removed by centrifugation at 15,000 rpm for 30 min at 4 °C. The clarified supernatant was incubated with Rabbit IgG Beads 4FF (SA082100, Smart Lifesciences) for 2 h at 4 °C with gentle rotation. The beads were washed with at least 5 column volumes of Wash buffer-1 (30 mM HEPES pH 7.4, 300 mM NaCl, 10% glycerol, 0.2 mM EDTA, 0.1% CHAPS, 5 mM MgCl_2_, 5 mM ATP, 2 mM DTT) followed by Wash buffer-2 (30 mM HEPES pH 7.4, 300 mM NaCl, 10% glycerol, 0.2 mM EDTA, 0.1% CHAPS, 5 mM MgCl_2_, 2 mM DTT). After washing, bound proteins were eluted by mixing the beads with an equal volume of 2xSDS loading buffer. Samples were separated on 4-20% SDS-PAGE gels at 250 V for 30 min, and analyzed by Coomassie Brilliant Blue staining or western blotting, as indicated (Anti-Strep II, AE066, Abclonal; anti-GAPDH, 60004-1-Ig, Proteintech).

### Human LAS1L-NOL9 complex purification

For LAS1L-NOL9 complex purification, full length LAS1L and LAS1L (H160A) were cloned into the pMLink vector with an N-terminal FLAG-4xProtein A-SUMO tag followed by TEV and 3C protease cleavage sites. NOL9 was cloned into pMLink vector without tag. 600 μg of each plasmid was transfected into 1 L of Expi293F cells (2 × 10^6^ cells mL^−1^) using PEI (40,000 MW, Yeasen). After 48 h, cells were harvested by centrifugation at 3,000 rpm for 10 min and resuspended in lysis buffer containing 30 mM HEPES (pH 7.4), 150 mM NaCl, 10% glycerol, 0.2 mM EDTA, 0.25% CHAPS, 5 mM MgCl_2_, 5 mM ATP, 2 mM DTT, and protease inhibitors (1 μg ml^−1^ aprotinin, 1 μg ml^−1^ pepstatin A, 1 μg ml^−1^ leupeptin, and 0.1 mM PMSF). Cell lysis was performed at 4 °C for 30 min with gentle rotation. Cell debris was removed by centrifugation at 15,000 rpm for 30 min at 4 °C. The clarified supernatant was incubated with Rabbit IgG Beads 4FF (SA082100, Smart Lifesciences) overnight at 4 °C with gentle rotation. The beads were washed with at least 5 column volumes of Wash buffer-1 containing 30 mM HEPES (pH 7.4), 500 mM NaCl, 10% glycerol, 0.2 mM EDTA, 0.1% CHAPS, 5 mM MgCl_2_, 5 mM ATP and 2 mM DTT, followed by Wash buffer-2 containing 30 mM HEPES (pH 7.4), 500 mM NaCl, 10% glycerol, 0.2 mM EDTA, 0.1% CHAPS, 5 mM MgCl_2_ and 2 mM DTT. The bound proteins were cleaved from the beads by incubating with Ulp1 protease at 4 °C overnight on a shaker at 200 rpm to prevent bead sedimentation. The cleaved proteins were eluted with Wash buffer-2, and fractions containing the LAS1L-NOL9 complex were pooled together. The pooled sample was diluted to reduce the NaCl concentration to 150 mM and then loaded onto a Mono Q 5/50 GL ion-exchange column (54807, Cytiva) pre-equilibrated with Buffer-A containing 30 mM HEPES (pH 7.4), 150 mM NaCl and 5 mM MgCl_2_. Proteins were eluted with a linear gradient of Buffer-B containing 30 mM HEPES (pH 7.4), 1 M NaCl and 5 mM MgCl_2_. Fractions containing the target complex were collected, concentrated using Amicon® Ultra Centrifugal Filter, 100 kDa MWCO (UFC9100, Millipore). The purified complex was used directly for cryo-EM grid preparation or flash-frozen in liquid nitrogen and stored at −80 °C.

### LAS1L-NOL9 endonuclease assays

A 27-nt RNA (5′-GUCGUUUUAGGUUUUACCAACUGCGGC-FAM-3′) based on yeast pre-ribosomal RNA was used as the model ITS2 substrate. The purified LAS1L-NOL9 complex was incubated with the RNA at the indicated concentrations for 1 h at 37 °C in a buffer containing 10 mM Tris-Cl (pH 8.0), 37.5 mM KCl, 2.5 mM ATP, 6.25 mM EDTA (pH 8.0), 4 mM DTT. After the reaction, samples were resolved on a 15% (v/v) denaturing urea polyacrylamide gel in 1x Tris-borate-EDTA buffer at 250 V for 30 min. The results were visualized using the Cy2 channel on an Amersham Typhoon 5 system (Cytiva).

### Generation of yeast strains

The TAP-Flag-Grc3 and the Rix1-FTpA strain were generated using standard yeast genetic methods^56,57^.Clones were selected on YEA plates supplemented with 100 µg/ml clonNAT and positive strains were confirmed by western blot and PCR.

### Grc3 and Rix1 sample preparation from *S*.*pombe*

Cells were grown in YPA medium until they reached an OD_600_ of about 2.5. The cells were then harvested by centrifugation and flash frozen in liquid nitrogen. A SPEX SamplePrep 6970EFM Freezer/Mill was used for cell lysis, and the cell powder was stored at −80 °C. The cell powder was then resuspended in a buffer solution containing 50 mM Tris-HCl (pH 7.5), 100 mM NaCl, 5 mM MgCl_2_, 1 mM DTT supplemented with 5% glycerol, 0.1% NP-40, and Complete EDTA-free protease inhibitor (Roche). The lysate was cleaned twice by centrifugation at 4 °C: first at 4,000 rpm for 15 minutes using an Eppendorf 5810R centrifuge, and second at 17,500 rpm for 25 minutes using a Sorvall LYNX 6000 Superspeed Centrifuge. IgG Sepharose 6 Fast Flow resin (Cytiva) was added to the cleared supernatant and incubated for 90 minutes at 4 °C under rotation. The IgG beads were collected by centrifugation at 1,800 rpm using an Eppendorf 5810 R centrifuge, washed once in a batch, and washed a second time using a 20-ml Econo-Pac chromatography column (Bio-Rad) via gravity flow. The IgG beads were resuspended in buffer supplemented with 5% glycerol, 0.1% NP-40, and homemade TEV protease, then incubated for 90 minutes at 16 °C under rotation. The supernatant was collected and incubated for 90 minutes with anti-FLAG^®^ M2 Affinity Gel (Sigma-Aldrich). The Flag beads were washed twice in batches, first with buffer supplemented with 0.01% NP-40, then with buffer supplemented with 0.05% octaethylene glycol monododecyl ether (Nikkol). One final wash was performed in a 1-ml gravity-flow column (Mobicol), and the samples were eluted with a buffer containing 0.05% octaethylene glycol monododecyl ether (Nikkol) and 3xFLAG peptide (Sigma-Aldrich, 250 μg/ml final concentration). Potential aggregates were removed by centrifugation in a tabletop centrifuge at 4 °C and 14,000 rpm for 5 minutes, after which the supernatant was used for cryo-EM grid preparation.

### Cryo-EM grid preparation and data acquisition

For the human LAS1L-NOL9 complex, the purified complex was applied to glow-discharged 300 mesh amorphous Ni/Ti foil (Au, 1.2/1.3) using a Vitrobot Mark IV (FEI Company) at 4 °C and 100% humidity. The sample was incubated on cryo-EM grids for 60 seconds (3.5 µl per grid) and then blotted for 3.5 seconds before being plunged into liquid ethane. Cryo-EM data were collected on a 300 keV Titan Krios G4 (FEI Company), equipped with a Falcon 4i detector and a Selectris X energy filter (Thermo Fisher Scientific) using EPU2 (Thermo Fisher Scientific). Collection was carried out with a nominal pixel size of 0.932Å/pixel, a defocus range of −1 to −2.5 µm and a total dose of 50 e^−^/Å^2^.

For the Grc3 and Rix1 samples from *S. pombe*, the purified samples were applied to glow-discharged R 3/3 holey copper grids with 3 nm continuous carbon support (Quantifoil) using a Vitrobot Mark IV (FEI Company) at 4 °C and 95% humidity. The samples were incubated on cryo-EM grids for 45 seconds (3.5 µl per grid) and then blotted for three seconds before being plunged into liquid ethane. Cryo-EM data were collected on a 300 keV Titan Krios G3 (FEI Company), equipped with a Falcon 4i detector and a Selectris X energy filter (Thermo Fisher Scientific) using EPU (Thermo Fisher Scientific). Collection was carried out with a nominal pixel size of 0.727 Å/pixel, a defocus range of −0.5 to −3.5 µm and a total dose of 40 e^−^/Å^2^.

For the human LAS1L-NOL9, *S. pombe* TAP-Flag-Grc3 and Rix1-FTpA datasets, 6,495, 46,644 and 64,904 movies were collected, respectively. MotionCor2^58^ was used for motion-correction of the movies and contrast-transfer function (CTF) parameters were estimated using CTFFIND4^59^.

### Electron microscopy and image processing

For the human LAS1L-NOL9 dataset, 2,261,561 particles were picked from the dataset using Gautomatch without a reference template. Particles were extracted with a pixel size of 1.864 Å/pixel and a box size of 150 × 150 pixels in Relion^60^, and then imported into cryoSPARC^61^ for 2D classification. A total of 499,721 good particles were selected and imported back to Relion to be extracted at 0.932 Å/pixel with a box size of 300 × 300 pixels. The resulting particles were then imported back to cryoSPARC^61^ for heterogeneous refinement and classified into four classes. All four classes displayed the classical butterfly-shaped architecture of the LAS1 complex; however, one class (C3) exhibited not only well-resolved density for LAS1L but also markedly better density for NOL9 compared to the other three classes. This class was therefore selected for the final reconstruction of the LAS1L-NOL9 complex. Particles from class C3 were subjected to non-uniform refinement to generate the final map, which was subsequently processed using local resolution filtering and DeepEMhancer^62^ to improve map interpretability.

For the Grc3 dataset, 44,647 micrographs were selected for data processing. The initial particles (3,614,528 particles) were picked using Relion (v4.0.1)^60^ autopick (Laplacian-of-Gaussian). The particles were extracted with a pixel size of 2.181 Å/pixel and a box size of 200 × 200 pixels, imported into cryoSPARC^61^ and used for 2D classification. Good 2D class averages were selected for the Rixosome (608,527 particles) and the pre-60S subunits (202,059 particles) and were used for ab-initio reconstructions and homogeneous refinements. The pre-60S particles were reimported into Relion and used for one round of 3D classification with six classes and without image alignment (see **Extended Data Fig. 6e**). To increase the number of Rixosome particles, particles were extracted with a pixel size of 1.454 Å/pixel and a box size of 240 × 240 pixels from patch CTF-estimated micrographs in cryoSPARC^61^. These particles were then used for two rounds of Topaz training, followed by heterogeneous refinement. The final Rixosome class contained 997,238 particles and was used for non-uniform refinement followed by 3D classification. All classes were extracted with a pixel size of 0.727 Å/pixel and a box size of 480 × 480 pixels, and were used for non-uniform refinements. While class C6 contained low-quality particles (188,027 particles; 6.85 Å), the other five classes were Rixosome states, with overall resolutions ranging from 2.76 Å to 3.02 Å. All classes contained the Las1 complex stably bound to one copy of the Rix1 complex. The second Rix1 complex exhibited conformational flexibility and was absent from the reconstitution of class C1, highly flexible in classes C2 and C3 and stably bound in classes C4 and C5. The single Rix1 Rixosome (class C1) was subjected to 2D classification and good 2D class averages were then used for non-uniform refinement (overall resolution of 2.74 Å). Class C5 had the most stable second Rix1 complex bound to the Las1 complex - double Rix1 Rixosome. Local refinements were performed for both classes after particle subtractions for the Las1 and the individual Rix1 complexes. For the Rix1 dataset, 60,247 micrographs were selected and used for data processing. Relion^60^ autopicking was used and 3,458,354 initial particles were extracted with a pixel size of 2.908 Å/pixel and a box size of 160 × 160 pixels. 2D classification was performed in cryoSPARC and 2D class averages of pre-60S particles were selected (1,323,582 particles) and used for ab-initio reconstruction and homogeneous refinement. Particles were imported into Relion and elaborated 3D classifications and focused 3D classifications were performed without image alignment (see sorting scheme **Extended Data Fig. 11d**). The final pre-60S states were extracted with a pixel size of 0.727 Å/pixel and a box size of 640 × 640 pixels and final non-uniform refinements and local refinements with particle subtraction were performed in cryoSPARC^61^ (see **Extended Data Figs. 12-14**). Composite maps of the local refinements were generated in ChimeraX^63^ with the vop max command.

For proximity pairing of pre-60S particles, particle coordinates were grouped by micrograph and matched using global minimum-distance assignment (Hungarian matching). Particle pairs with center-to-center distances above 350 Å were discarded. For the remaining pairs, the midpoint between the coordinates of each pre-60S in a pair was used as the new extraction coordinates. Paired and centered particles were re-extracted with a pixel size of 2.908 Å/pixel (box size 230 × 230 pixels) and subject to 2D classification in cryoSPARC (**Fig. 6e**).

### Model building and refinement

For model building of the human LAS1L-NOL9 complex, initial complex model of LAS1L and NOL9 were generated using AlphaFold3^64^. These predicted models were rigid-body fitted into the cryo-EM density map using ChimeraX^63^. The fitted models were then manually adjusted and rebuilt residue-by-residue in Coot^65^ to improve agreement with the density. Regions lacking clear density (e.g., HD) were not modeled.

The molecular models of the Rixosome were built using the following structures as starting points: the Las1 complex from *C. thermophilum* (PDB: 6OF2, 8PUW)^7,41^, the Rix1 complex models from *S. cerevisiae* (PDB ID: 6YLE)^19^ and *C. thermophilum* (PDB ID: 8PTW)^7^, as well as AlphaFold and AlphaFold multimer predictions of the *S. pombe* components. All models were rigid-body fitted into the cryo-EM densities using ChimeraX^63^ and were used to build the *S*.*pombe* Rixosome complexes in Coot (v0.9.8.96)^65^. For the Las1 HD-Ipi3 CC model, the AlphaFold multimer prediction was fitted into the cryo-EM density and manually adjusted with Coot^65^.

To generate the molecular models of the *S. pombe* pre-60S states, we started by building state A1 and B3 in Coot^65^. For both states, we used the *S*.*pombe* 80S ribosome (PDB ID: 9AXU)^66^ and the nucleolar pre-60S from *S*.*pombe* (PDB ID: 8ESQ)^67^ as initial models. Additionally, we used the models of pre-60S states from *S. cerevisiae* (PDB: 7UOO)^68^ and *C. thermophilum* (PDB ID: 8PV7)^7^ for state A1, as well as the *S. cerevisiae* (PDB ID: 6YLH)^19^ and *C. thermophilum* (PDB ID: 8PV6)^7^ models for state B3. Alphafold prediction of the individual protein chains were used to further assist model building. For state A2, the state A1 was used and the ES27 was built manually in Coot. For models B1, B2, C1, C2, C3 and C4 the state B3 was used and parts of the model missing in the individual states were removed and the models were adjusted in Coot.

The final models were real-space refined with Phenix (v2.1-6048)^69^ with secondary structure restraints applied. The final model was validated using MolProbity^70^. Structure figures were generated with ChimeraX (v1.9)^63^.

## Acknowledgements

We thank Susanne Rieder and Charlotte Ungewickell, and the Core Facility of Shanghai Medical College, Fudan University and Center of Cryo-Electron Microscopy at Fudan University for technical support.

## Author Contributions

Conceptualization, M.T., J.Z., R.B. and J.C. Methodology, M.T., J.Z., X.H., T.D., J.S., S.N., O.B., Y.L. Investigation, M.T., J.Z., X.H., T.D., J.S., S.N., J.L. Visualization, M.T., X.H. and J.C. Project administration, M.T., R.B., J.C. Supervision, M.T. and J.C. Funding acquisition, R.B. and J.C. Writing— original draft, M.T., J.Z., X.H. and J.C. Writing— review, and editing, all authors

## Data availability

The final cryo-EM maps/composite maps and molecular models were deposited to the Electron Microscopy Data Bank (EMDB) and the Protein Data Bank (PDB): Human LAS1L-NOL9 complex (EMD-XXXXX, XXXX), single Rix1 Rixosome (EMD-XXXXX, XXXX), double Rix1 Rixosome (EMD-XXXXX, XXXX), pre-60S State A1 (EMD-XXXXX, XXXX), State A2 (EMD-XXXXX, XXXX), State B1 (EMD-XXXXX, XXXX), State B2 (EMD-XXXXX, XXXX), State B3 (EMD-XXXXX, XXXX), State C1 (EMD-XXXXX, XXXX), State C2 (EMD-XXXXX, XXXX), State C3 (EMD-XXXXX, XXXX), State C4 (EMD-XXXXX, XXXX). The consensus maps and all local refinement maps used for the composite maps were deposited to the Electron Microscopy Data Bank (EMDB) and are listed in **Extended Data Tables 2** and **3**.

## References

1 Vanden Broeck, A. & Klinge, S. Eukaryotic Ribosome Assembly. Annual review of biochemistry 93, 189–210, doi:10.1146/annurev-biochem-030222-113611 (2024).

2 Hurt, E., Iwasa, J. & Beckmann, R. SnapShot: Eukaryotic ribosome biogenesis II. Cell 187, 1314–1314 e1311, doi:10.1016/j.cell.2024.01.043 (2024).

3 Bassler, J. & Hurt, E. Eukaryotic Ribosome Assembly. Annual review of biochemistry 88, 281–306, doi:10.1146/annurev-biochem-013118-110817 (2019).

4 Woolford, J. L., Jr. & Baserga, S. J. Ribosome biogenesis in the yeast Saccharomyces cerevisiae. Genetics 195, 643–681, doi:10.1534/genetics.113.153197 (2013).

5 Vanden Broeck, A. & Klinge, S. Principles of human pre-60S biogenesis. Science 381, eadh3892, doi:10.1126/science.adh3892 (2023).

6 Kater, L. et al. Visualizing the Assembly Pathway of Nucleolar Pre-60S Ribosomes. Cell 171, 1599–1610 e1514, doi:10.1016/j.cell.2017.11.039 (2017).

7 Thoms, M. et al. Structural insights into coordinating 5S RNP rotation with ITS2 pre-RNA processing during ribosome formation. EMBO reports 24, e57984, doi:10.15252/embr.202357984 (2023).

8 Sanghai, Z. A. et al. Modular assembly of the nucleolar pre-60S ribosomal subunit. Nature 556, 126–129, doi:10.1038/nature26156 (2018).

9 Zhou, D. et al. Cryo-EM structure of an early precursor of large ribosomal subunit reveals a half-assembled intermediate. Protein Cell 10, 120–130, doi:10.1007/s13238-018-0526-7 (2019).

10 Granneman, S., Petfalski, E. & Tollervey, D. A cluster of ribosome synthesis factors regulate pre-rRNA folding and 5.8S rRNA maturation by the Rat1 exonuclease. The EMBO journal 30, 4006–4019, doi:10.1038/emboj.2011.256 (2011).

11 Zhang, Y. et al. Visualizing the nucleoplasmic maturation of human pre-60S ribosomal particles. Cell research 33, 867–878, doi:10.1038/s41422-023-00853-9 (2023).

12 Lau, B. et al. Mechanism of 5S RNP recruitment and helicase-surveilled rRNA maturation during pre-60S biogenesis. EMBO reports 24, e56910, doi:10.15252/embr.202356910 (2023).

13 Sanghai, Z. A., Piwowarczyk, R., Broeck, A. V. & Klinge, S. A co-transcriptional ribosome assembly checkpoint controls nascent large ribosomal subunit maturation. Nature structural & molecular biology 30, 594–599, doi:10.1038/s41594-023-00947-3 (2023).

14 Bassler, J. et al. The AAA-ATPase Rea1 drives removal of biogenesis factors during multiple stages of 60S ribosome assembly. Molecular cell 38, 712–721, doi:10.1016/j.molcel.2010.05.024 (2010).

15 Mitterer, V. et al. Concurrent remodelling of nucleolar 60S subunit precursors by the Rea1 ATPase and Spb4 RNA helicase. eLife 12, doi:10.7554/eLife.84877 (2023).

16 Ahmed, Y. L., Thoms, M., Mitterer, V., Sinning, I. & Hurt, E. Crystal structures of Rea1-MIDAS bound to its ribosome assembly factor ligands resembling integrin-ligand-type complexes. Nature communications 10, 3050, doi:10.1038/s41467-019-10922-6 (2019).

17 Thoms, M., Ahmed, Y. L., Maddi, K., Hurt, E. & Sinning, I. Concerted removal of the Erb1-Ytm1 complex in ribosome biogenesis relies on an elaborate interface. Nucleic acids research 44, 926–939, doi:10.1093/nar/gkv1365 (2016).

18 Wegrecki, M., Rodriguez-Galan, O., de la Cruz, J. & Bravo, J. The structure of Erb1-Ytm1 complex reveals the functional importance of a high-affinity binding between two beta-propellers during the assembly of large ribosomal subunits in eukaryotes. Nucleic acids research 43, 11017–11030, doi:10.1093/nar/gkv1043 (2015).

19 Kater, L. et al. Construction of the Central Protuberance and L1 Stalk during 60S Subunit Biogenesis. Molecular cell 79, 615–628 e615, doi:10.1016/j.molcel.2020.06.032 (2020).

20 Schuller, J. M., Falk, S., Fromm, L., Hurt, E. & Conti, E. Structure of the nuclear exosome captured on a maturing preribosome. Science 360, 219–222, doi:10.1126/science.aar5428 (2018).

21 Gasse, L., Flemming, D. & Hurt, E. Coordinated Ribosomal ITS2 RNA Processing by the Las1 Complex Integrating Endonuclease, Polynucleotide Kinase, and Exonuclease Activities. Molecular cell 60, 808–815, doi:10.1016/j.molcel.2015.10.021 (2015).

22 Castle, C. D., Cassimere, E. K. & Denicourt, C. LAS1L interacts with the mammalian Rix1 complex to regulate ribosome biogenesis. Mol Biol Cell 23, 716–728, doi:10.1091/mbc.E11-06-0530 (2012).

23 Pillon, M. C., Gordon, J., Frazier, M. N. & Stanley, R. E. HEPN RNases - an emerging class of functionally distinct RNA processing and degradation enzymes. Critical reviews in biochemistry and molecular biology 56, 88–108, doi:10.1080/10409238.2020.1856769 (2021).

24 Pillon, M. C. et al. It takes two (Las1 HEPN endoribonuclease domains) to cut RNA correctly. The Journal of biological chemistry 295, 5857–5870, doi:10.1074/jbc.RA119.011193 (2020).

25 Mitchell, P., Petfalski, E., Shevchenko, A., Mann, M. & Tollervey, D. The exosome: a conserved eukaryotic RNA processing complex containing multiple 3’-->5’ exoribonucleases. Cell 91, 457–466, doi:10.1016/s0092-8674(00)80432-8 (1997).

26 Thoms, M. et al. The Exosome Is Recruited to RNA Substrates through Specific Adaptor Proteins. Cell 162, 1029–1038, doi:10.1016/j.cell.2015.07.060 (2015).

27 Falk, S. et al. Structural insights into the interaction of the nuclear exosome helicase Mtr4 with the preribosomal protein Nop53. Rna 23, 1780–1787, doi:10.1261/rna.062901.117 (2017).

28 Fromm, L. et al. Reconstitution of the complete pathway of ITS2 processing at the pre-ribosome. Nature communications 8, 1787, doi:10.1038/s41467-017-01786-9 (2017).

29 Thomson, E. & Tollervey, D. The final step in 5.8S rRNA processing is cytoplasmic in Saccharomyces cerevisiae. Molecular and cellular biology 30, 976–984, doi:10.1128/MCB.01359-09 (2010).

30 Gordon, J. et al. PELP1 coordinates the modular assembly and enzymatic activity of the rixosome complex. Sci Adv 11, eadw4603, doi:10.1126/sciadv.adw4603 (2025).

31 Barrio-Garcia, C. et al. Architecture of the Rix1-Rea1 checkpoint machinery during pre-60S-ribosome remodeling. Nature structural & molecular biology 23, 37–44, doi:10.1038/nsmb.3132 (2016).

32 Sarkar, A. et al. Preribosomes escaping from the nucleus are caught during translation by cytoplasmic quality control. Nature structural & molecular biology 24, 1107–1115, doi:10.1038/nsmb.3495 (2017).

33 Rodriguez-Galan, O., Garcia-Gomez, J. J., Kressler, D. & de la Cruz, J. Immature large ribosomal subunits containing the 7S pre-rRNA can engage in translation in Saccharomyces cerevisiae. RNA biology 12, 838–846, doi:10.1080/15476286.2015.1058477 (2015).

34 Biedka, S. et al. Hierarchical recruitment of ribosomal proteins and assembly factors remodels nucleolar pre-60S ribosomes. The Journal of cell biology 217, 2503–2518, doi:10.1083/jcb.201711037 (2018).

35 Butterfield, R. J. et al. Congenital lethal motor neuron disease with a novel defect in ribosome biogenesis. Neurology 82, 1322–1330, doi:10.1212/WNL.0000000000000305 (2014).

36 Hu, H. et al. X-exome sequencing of 405 unresolved families identifies seven novel intellectual disability genes. Mol Psychiatry 21, 133–148, doi:10.1038/mp.2014.193 (2016).

37 Zhou, H. et al. Rixosomal RNA degradation contributes to silencing of Polycomb target genes. Nature 604, 167–174, doi:10.1038/s41586-022-04598-0 (2022).

38 Shipkovenska, G., Durango, A., Kalocsay, M., Gygi, S. P. & Moazed, D. A conserved RNA degradation complex required for spreading and epigenetic inheritance of heterochromatin. eLife 9, doi:10.7554/eLife.54341 (2020).

39 Huang, J. & Tong, L. Molecular insights into the overall architecture of human rixosome. Nature communications 16, 3288, doi:10.1038/s41467-025-58732-3 (2025).

40 Chen, J. et al. Structural and mechanistic insights into ribosomal ITS2 RNA processing by nuclease-kinase machinery. Elife 12, doi:10.7554/eLife.86847 (2024).

41 Pillon, M. C. et al. Cryo-EM reveals active site coordination within a multienzyme pre-rRNA processing complex. Nat Struct Mol Biol 26, 830–839, doi:10.1038/s41594-019-0289-8 (2019).

42 Micic, J. et al. Coupling of 5S RNP rotation with maturation of functional centers during large ribosomal subunit assembly. Nature communications 11, 3751, doi:10.1038/s41467-020-17534-5 (2020).

43 Wu, S. et al. Diverse roles of assembly factors revealed by structures of late nuclear pre-60S ribosomes. Nature 534, 133–137, doi:10.1038/nature17942 (2016).

44 Leidig, C. et al. 60S ribosome biogenesis requires rotation of the 5S ribonucleoprotein particle. Nature communications 5, 3491, doi:10.1038/ncomms4491 (2014).

45 Fiorentino, F. et al. Highly conserved ribosome biogenesis pathways between human and yeast revealed by the MDN1-NLE1 interaction and NLE1 containing pre-60S subunits. Nucleic Acids Res 53, doi:10.1093/nar/gkaf255 (2025).

46 Cruz, V. E. et al. Sequence-specific remodeling of a topologically complex RNP substrate by Spb4. Nature structural & molecular biology 29, 1228–1238, doi:10.1038/s41594-022-00874-9 (2022).

47 Cruz, V. E., Weirich, C. S., Peddada, N. & Erzberger, J. P. The DEAD-box ATPase Dbp10/DDX54 initiates peptidyl transferase center formation during 60S ribosome biogenesis. Nature communications 15, 3296, doi:10.1038/s41467-024-47616-7 (2024).

48 Pillon, M. C. & Stanley, R. E. Nuclease integrated kinase super assemblies (NiKs) and their role in RNA processing. Current genetics 64, 183–190, doi:10.1007/s00294-017-0749-9 (2018).

49 Jumper, J. et al. Highly accurate protein structure prediction with AlphaFold. Nature 596, 583–589, doi:10.1038/s41586-021-03819-2 (2021).

50 Gordon, J. et al. Cryo-EM reveals the architecture of the PELP1-WDR18 molecular scaffold. Nat Commun 13, 6783, doi:10.1038/s41467-022-34610-0 (2022).

51 Pillon, M. C., Sobhany, M. & Stanley, R. E. Characterization of the molecular crosstalk within the essential Grc3/Las1 pre-rRNA processing complex. Rna 24, 721–738, doi:10.1261/rna.065037.117 (2018).

52 Pillon, M. C., Sobhany, M., Borgnia, M. J., Williams, J. G. & Stanley, R. E. Grc3 programs the essential endoribonuclease Las1 for specific RNA cleavage. Proceedings of the National Academy of Sciences of the United States of America 114, E5530–E5538, doi:10.1073/pnas.1703133114 (2017).

53 Ulbrich, C. et al. Mechanochemical removal of ribosome biogenesis factors from nascent 60S ribosomal subunits. Cell 138, 911–922, doi:10.1016/j.cell.2009.06.045 (2009).

54 Zhao, X. et al. In-cell structures visualize human pre-ribosome assembly in the nucleolus (openRxiv, 2026).

55 Lu, P. et al. Three-dimensional structure of human gamma-secretase. Nature 512, 166–170, doi:10.1038/nature13567 (2014).

56 Janke, C. et al. A versatile toolbox for PCR-based tagging of yeast genes: new fluorescent proteins, more markers and promoter substitution cassettes. Yeast 21, 947–962, doi:10.1002/yea.1142 (2004).

57 Longtine, M. S. et al. Additional modules for versatile and economical PCR-based gene deletion and modification in Saccharomyces cerevisiae. Yeast 14, 953–961, doi:10.1002/(SICI)1097-0061(199807)14:10<953::AID-YEA293>3.0.CO;2-U (1998).

58 Zheng, S. Q. et al. MotionCor2: anisotropic correction of beam-induced motion for improved cryo-electron microscopy. Nat Methods 14, 331–332, doi:10.1038/nmeth.4193 (2017).

59 Rohou, A. & Grigorieff, N. CTFFIND4: Fast and accurate defocus estimation from electron micrographs. J Struct Biol 192, 216–221, doi:10.1016/j.jsb.2015.08.008 (2015).

60 Kimanius, D., Dong, L., Sharov, G., Nakane, T. & Scheres, S. H. W. New tools for automated cryo-EM single-particle analysis in RELION-4.0. Biochem J 478, 4169–4185, doi:10.1042/BCJ20210708 (2021).

61 Punjani, A., Rubinstein, J. L., Fleet, D. J. & Brubaker, M. A. cryoSPARC: algorithms for rapid unsupervised cryo-EM structure determination. Nat Methods 14, 290–296, doi:10.1038/nmeth.4169 (2017).

62 Sanchez-Garcia, R. et al. DeepEMhancer: a deep learning solution for cryo-EM volume post-processing. Commun Biol 4, 874, doi:10.1038/s42003-021-02399-1 (2021).

63 Meng, E. C. et al. UCSF ChimeraX: Tools for structure building and analysis. Protein Sci 32, e4792, doi:10.1002/pro.4792 (2023).

64 Abramson, J. et al. Accurate structure prediction of biomolecular interactions with AlphaFold 3. Nature 630, 493–500, doi:10.1038/s41586-024-07487-w (2024).

65 Emsley, P. & Cowtan, K. Coot: model-building tools for molecular graphics. Acta Crystallogr D Biol Crystallogr 60, 2126–2132, doi:10.1107/S0907444904019158 (2004).

66 Gemin, O. et al. Ribosomes hibernate on mitochondria during cellular stress. Nature communications 15, 8666, doi:10.1038/s41467-024-52911-4 (2024).

67 Ugolini, I. et al. Chromatin localization of nucleophosmin organizes ribosome biogenesis. Molecular cell 82, 4443–4457 e4449, doi:10.1016/j.molcel.2022.10.033 (2022).

68 Sekulski, K., Cruz, V. E., Weirich, C. S. & Erzberger, J. P. rRNA methylation by Spb1 regulates the GTPase activity of Nog2 during 60S ribosomal subunit assembly. Nat Commun 14, 1207, doi:10.1038/s41467-023-36867-5 (2023).

69 Liebschner, D. et al. Macromolecular structure determination using X-rays, neutrons and electrons: recent developments in Phenix. Acta Crystallogr D Struct Biol 75, 861–877, doi:10.1107/S2059798319011471 (2019).

70 Chen, V. B. et al. MolProbity: all-atom structure validation for macromolecular crystallography. Acta crystallographica. Section D, Biological crystallography 66, 12–21, doi:10.1107/S0907444909042073 (2010).

