## Extended Data and Tables for "Dynamic architecture of the Rixosome reveals mechanism of activation and ITS2 processing"

**Extended Data associated with this article includes:**
Extended Data Figure 1-16
Extended Data Table 1-3

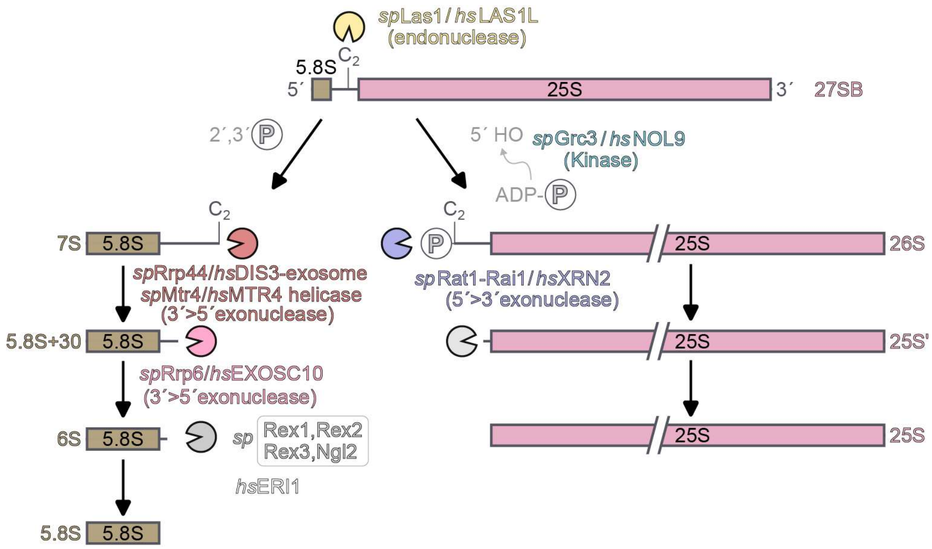

**Extended Data Fig. 1 | General schematic overview of ITS2 processing during yeast pre-60S maturation.** Schematic representation of the Las1-mediated $C_2$ cleavage within the ITS2 and subsequent processing of the 5.8S and 25S rRNA precursors. Cleavage at the $C_2$ site generates the 7S pre-rRNA and the 26S pre-rRNA, the precursors of the 5.8S rRNA and 25S rRNA, respectively. The cleaved 26S pre-rRNA contains a 5'-OH end, which is phosphorylated by Grc3 to allow Rat1-Rai1 (also named Dhp1-Din1 in *S. pombe*) mediated 5' to 3' trimming of the precursor. The 7S pre-rRNA harbors a 2'3' cyclic phosphate at the 3' end, and processing occurs through a successive pathway starting with nuclear exosome and the RNA helicase Mtr4, generating the 5.8S+30, which is further trimmed by Rrp6 and downstream nucleases to generate mature 5.8S rRNA.

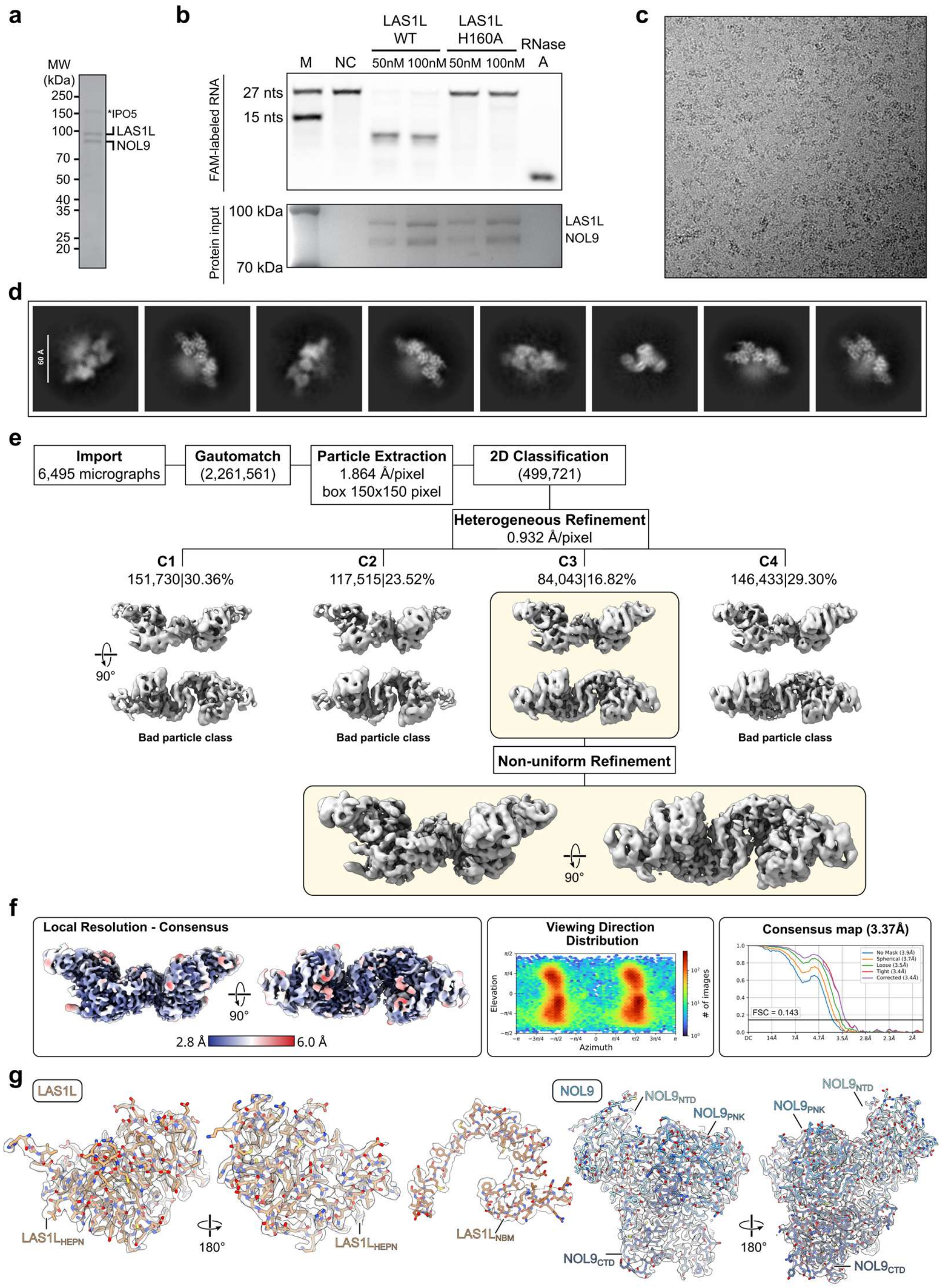

**Extended Data Fig. 2 | Biochemical validation and cryo-EM processing of the human LAS1 complex. a**, SDS PAGE analysis of the purified human LAS1 complex. **b**, *In vitro* ITS2 cleavage assay using the purified wild type LAS1 complex and the catalytic inactive LAS1L H160A complex. RNase A was used as a positive control. Yeast ITS2 fragment containing the $C_2$ cleavage site was used as substrate. **c-d**, Representative cryo-EM micrograph (**c**) and representative 2D class averages (**d**) of the human LAS1 complex. **e**, Cryo-EM data processing workflow of the human LAS1 complex dataset. **f**, Local resolution map, orientation distribution, and Fourier shell correlation curves of the final reconstruction. **g**, Representative cryo-EM density maps and fitted atomic models of LAS1L and NOL9.

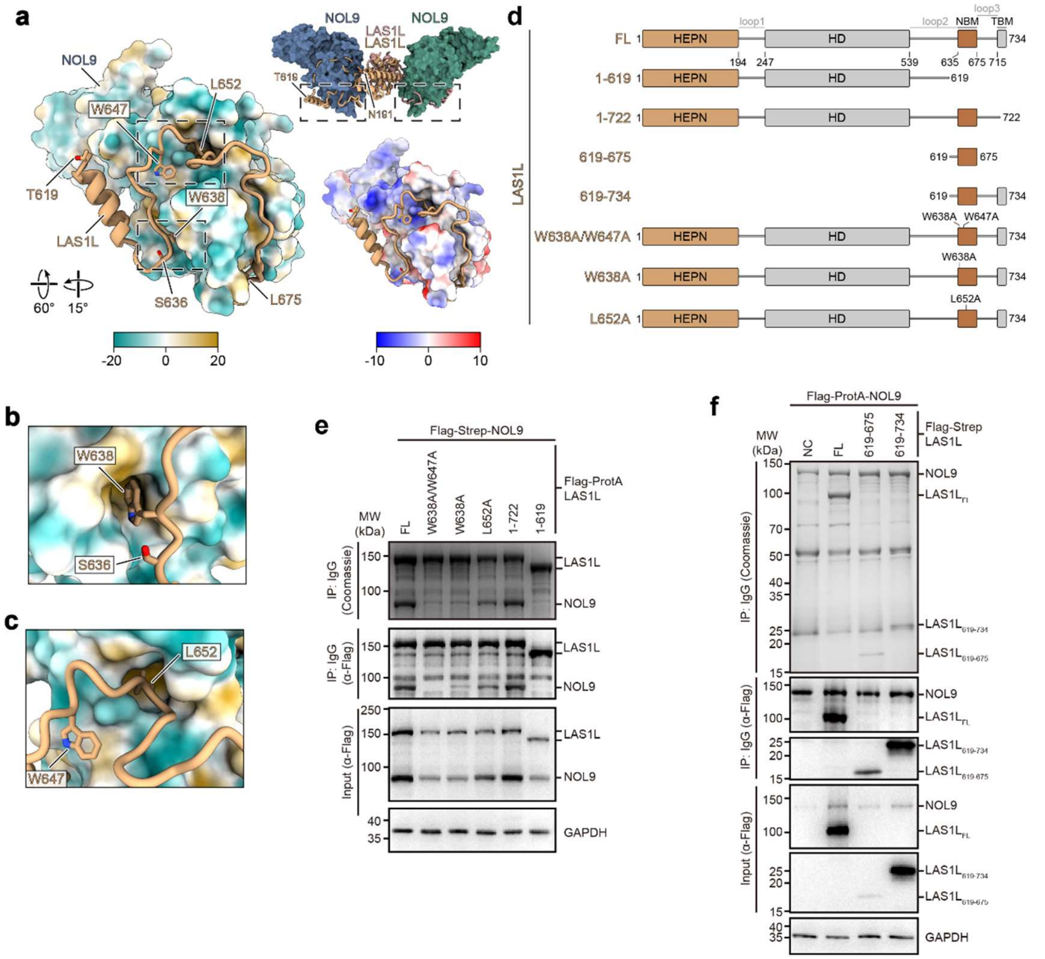

**Extended Data Fig. 3 | LAS1L NBM mediates assembly of the human LAS1 complex. a,** Overall position of the LAS1L NBM in the human LAS1 complex (top right). The lower panels show the NOL9 surface, colored according to lipophilicity (left) and electrostatic surface potential (right). The LAS1L NBM binds within a broad surface groove of NOL9. **b-c**, Close up views of the interactions mediated by the LAS1L residues W638, W647 and L652. **d**, Schematic representation of LAS1L truncations and mutants used for mapping the NOL9 binding region. **e**, Affinity purification analysis of the interaction between Flag-ProtA-LAS1L variants and Flag-Strep-NOL9. **f**, Affinity purification analysis showing that the LAS1L fragment containing the NBM (aa. 619-675) is sufficient for NOL9 binding.

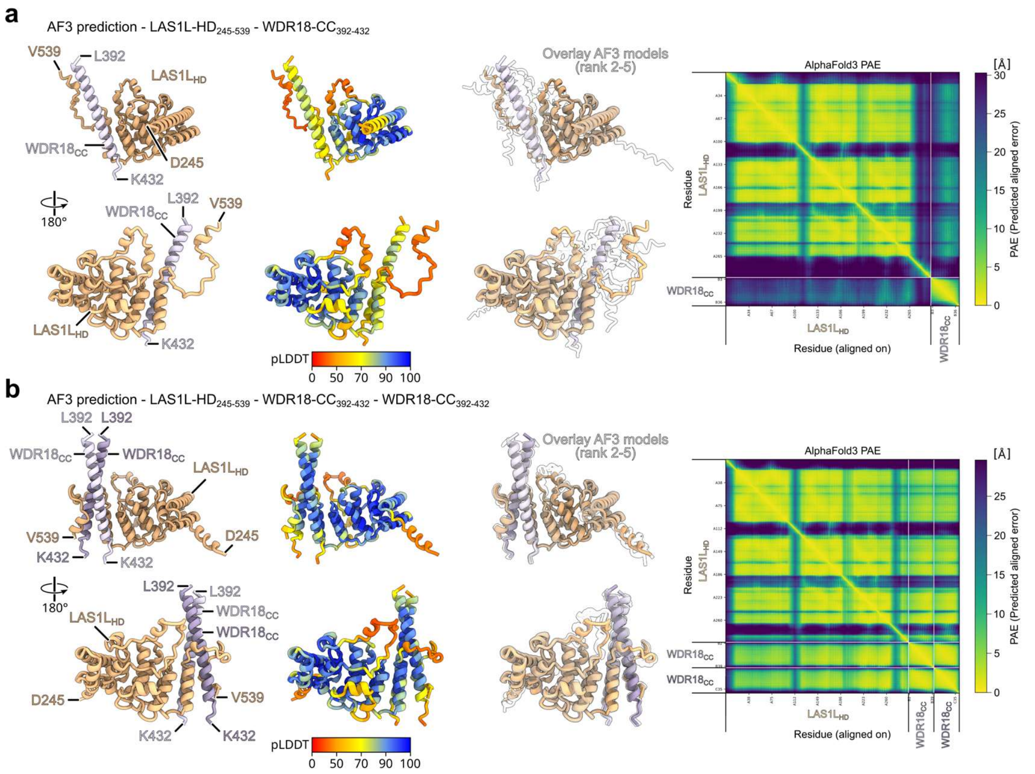

**Extended Data Fig. 4 | AlphaFold3 prediction of the LAS1L HD and WDR18 CC interaction.** **a-b,** AlphaFold3 predictions of the interaction between the human LAS1L HD and the C-terminal coiled-coil helix of WDR18 (CC, aa. 392-432). Models are shown for human LAS1L HD bound to one WDR18 CC helix (**a**) or to the WDR18 CC dimer (**b**). For each prediction, the top ranked model, pLDDT colored model, the overlay of ranked models 2-5, and the predicted aligned error (PAE) plot are shown.

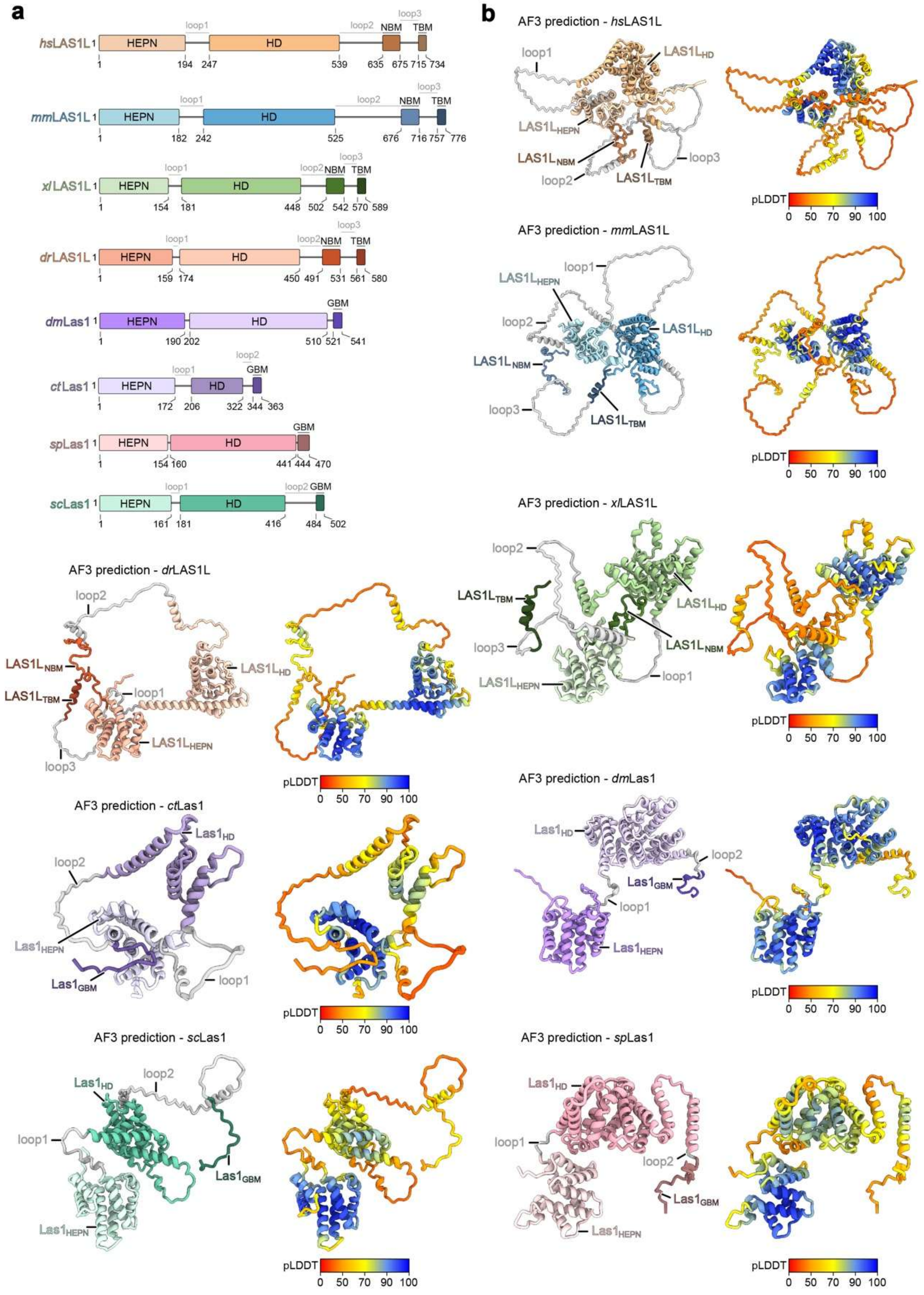

**Extended Data Fig. 5 | Comparison of Las1 domain architectures across species. a-b,** Domain architecture (**a**) and AlphaFold3 predicted structural models (**b**) of Las1 homologs from human (*hs*), mouse (*mm*), frog (*xl*), zebrafish (*dr*), fruit fly (*dm*), *Chaetomium thermophilum* (*ct*), *S. pombe* (*sp*) and *Saccharomyces cerevisiae* (*sc*). The *S. pombe* Las1 contains shortened loop regions flanking the HD. The connecting loop regions within the Las1 homologs are colored gray, and the predicted models are also colored according to their pLDDT scores.

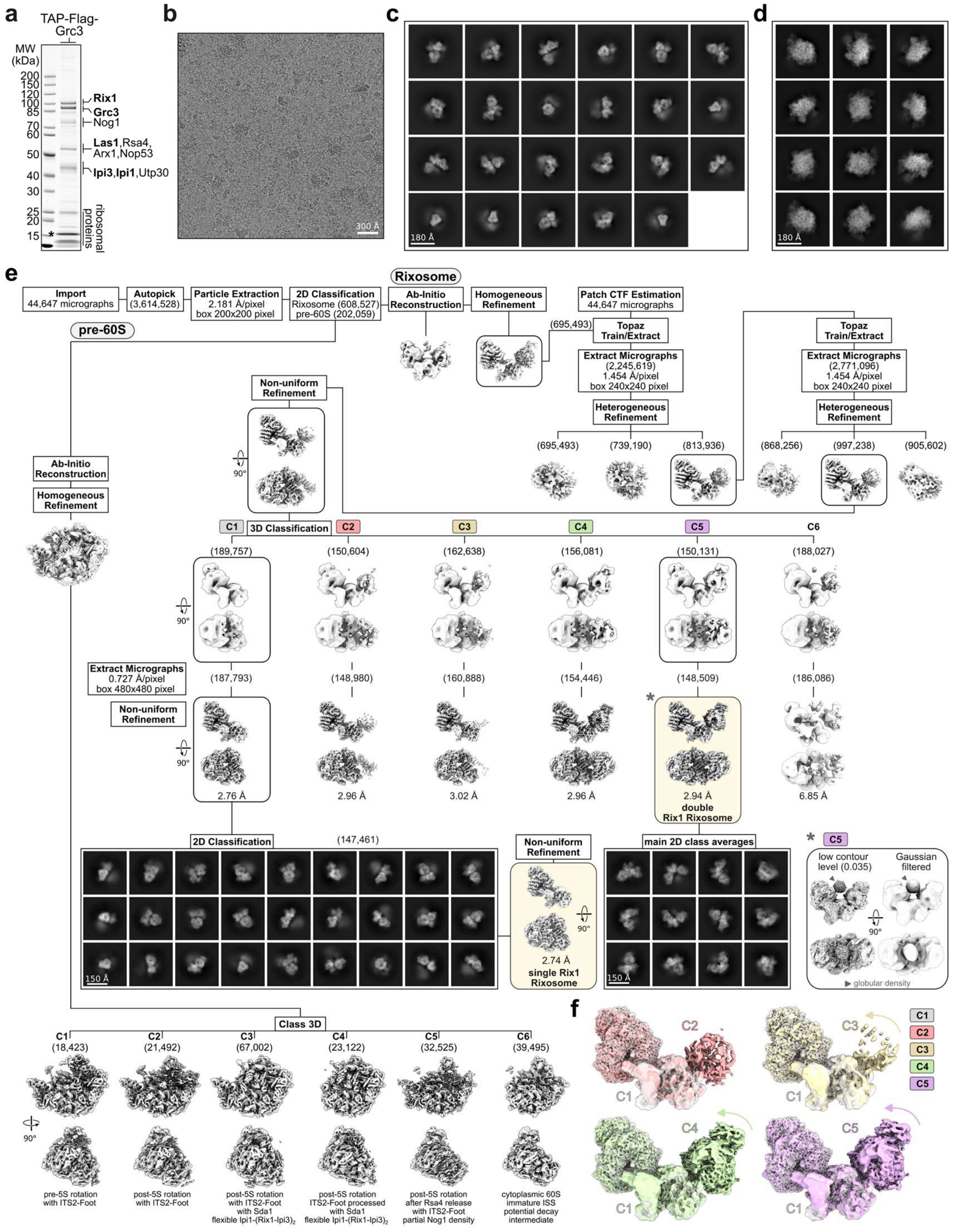

**Extended Data Fig. 6 | Cryo-EM analysis of the Grc3 dataset. a**, Coomassie stained SDS-PAGE of the Grc3 purification. The asterisk marks the contamination of Cam1. **b**, Representative cryo-EM micrograph of the Grc3 dataset. **c-d**, 2D class averages of the Rixosome (**c**) and pre-60S classes (**d**). **e**, Cryo-EM sorting scheme of the Grc3 dataset. Particle numbers are shown in brackets. The double Rix1 Rixosome (class C5, *) is also shown at a low contour level and Gaussian filtered to highlight the flexible, global density between the two Rix1 complex copies (marked with an arrow). **f**, Comparison of the conformational states of the Rixosome obtained through 3D classification (see **e**). The Las1 complex and one Rix1 complex form a rigid assembly in all observed classes. In contrast, the second Rix1 complex exhibits conformational flexibility, being flexible attached in classes C2 and C3, and more stable associated in classes C4 and C5.

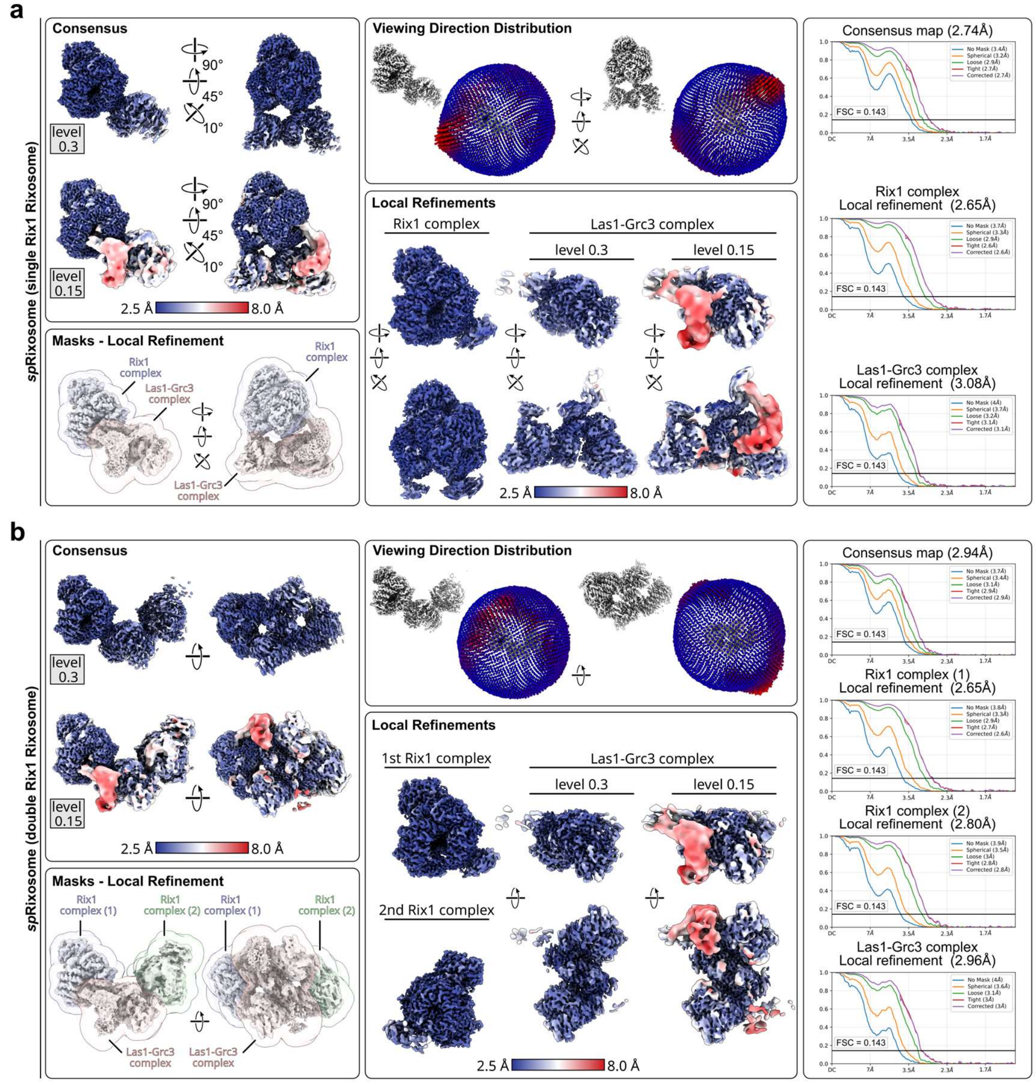

**Extended Data Fig. 7 | Map validation of the *S. pombe* Rixosome structures. a-b**, Cryo-EM validation of the single Rix1 Rixosome (**a**) and the double Rix1 Rixosome (**b**) reconstructions, including local resolution maps, orientation distributions, mask based local refinement strategies, and Fourier shell correlation (FSC) curves, are shown. Overall maps and Las1 focused maps are also shown at low contour levels to visualize the lower resolution density of the Las1 HD region.

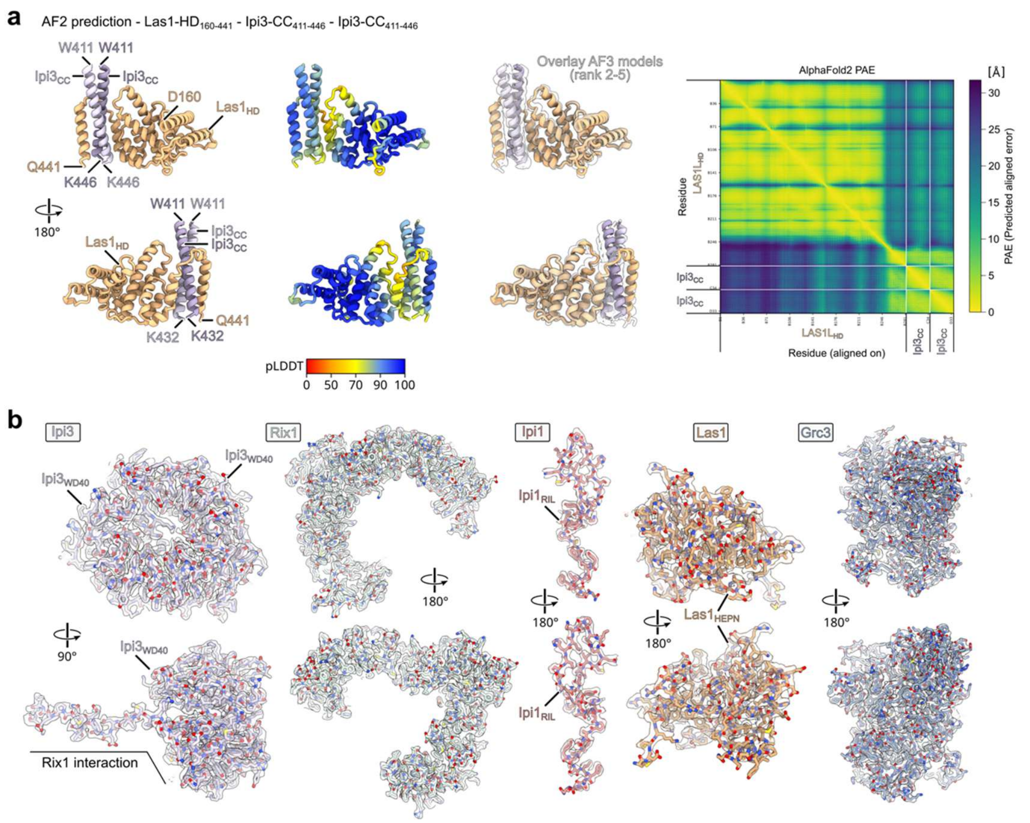

**Extended Data Fig. 8 | Model validation of the *S. pombe* Rixosome structure. a**, AlphaFold2 multimer prediction of the interaction between Las1 HD (aa. 160-441) and the Ipi3 CC dimer (aa. 411-446). The predicted model, pLDDT colored model, overlays with ranked models 2-5 and the predicted aligned error (PAE) plot are shown. **b**, Representative cryo-EM densities and fitted atomic models of Ipi3, Rix1, Ipi1, Las1 and Grc3 in the *S. pombe* Rixosome structure, showing good agreement between the models and the cryo-EM densities.

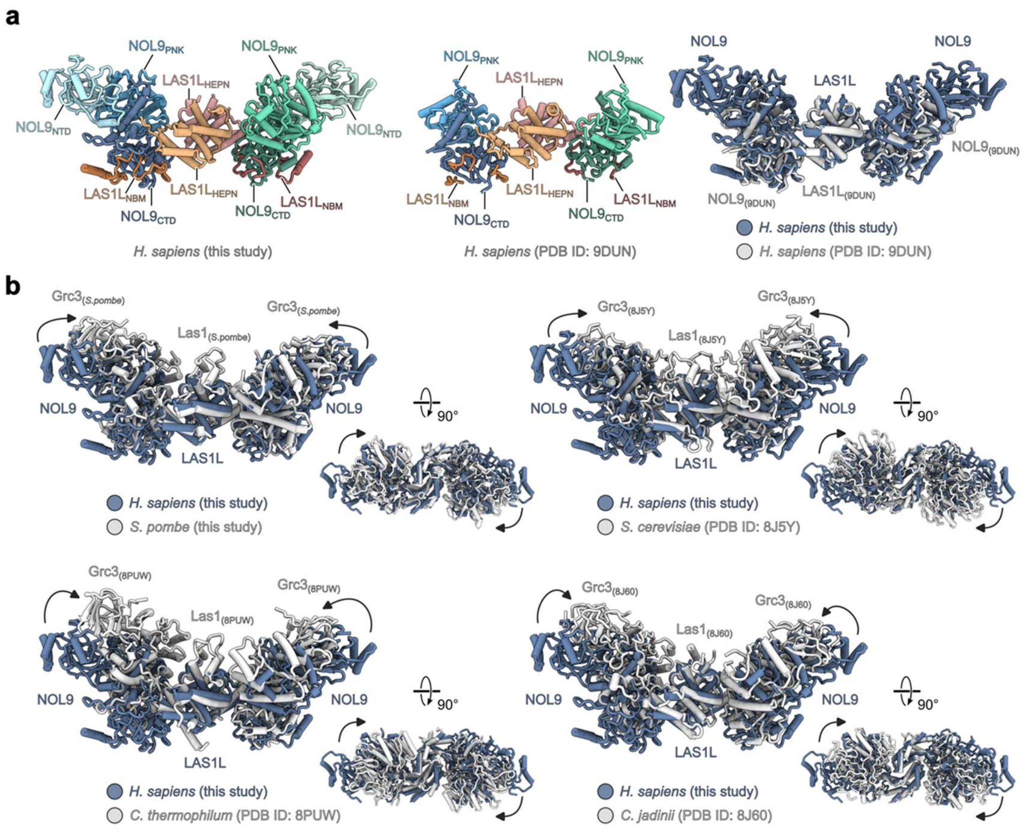

**Extended Data Fig. 9 | Structural comparison of different Las1 complexes. a**, Structural comparison of the human LAS1 complex determined in this study with the previously reported human LAS1 complex (PDB ID: 9DUN). The comparison is shown in the same orientation, highlighting the improved molecular model of the present LAS1 complex structure. **b**, Structural comparison of the human LAS1 complex (dark blue) with fungal Las1 complexes (gray) from *S. pombe* (this study), *S. cerevisiae* (PDB ID: 8J5Y), *C. thermophilum* (PDB ID: 8PUW), and *C. jadinii* (PDB ID: 8J60). The human LAS1 complex adopts a more open conformation compared to the fungal Las1 complexes.

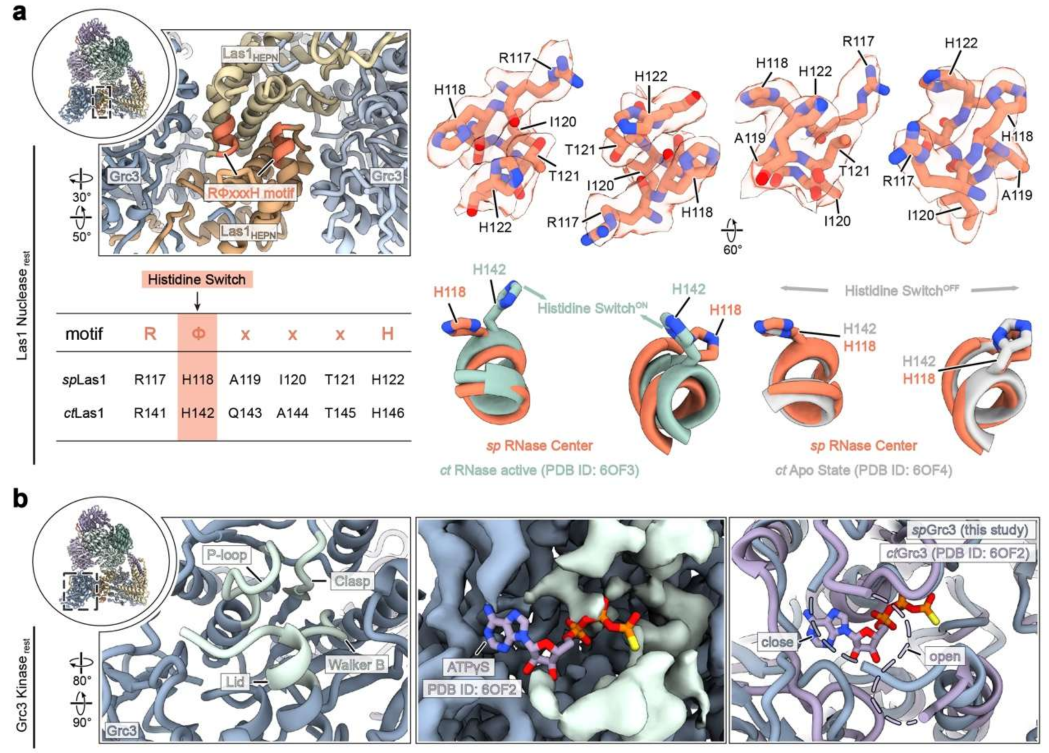

**Extended Data Fig. 10 | Catalytic states of the Las1 and Grc3 active centers. a**, Structure of the Las1 nuclease center (left). The conserved histidine switch motif is shown, together with the corresponding cryo-EM density, in two different views. Structural comparison with active (PDB ID: 6OF3) and the resting apo (PDB ID: 6OF4) state Las1 nuclease centers shows that Las1 adopts a resting conformation in the free Rixosome. **b**, Structure of the Grc3 kinase center. The P loop, lid, and Walker B motif are indicated. Structural comparison with the ATPγS bound Grc3 structure (PDB ID: 6OF2) from *C. thermophilum* shows that the Grc3 kinase center adopts a closed, nucleotide free conformation in the free Rixosome.

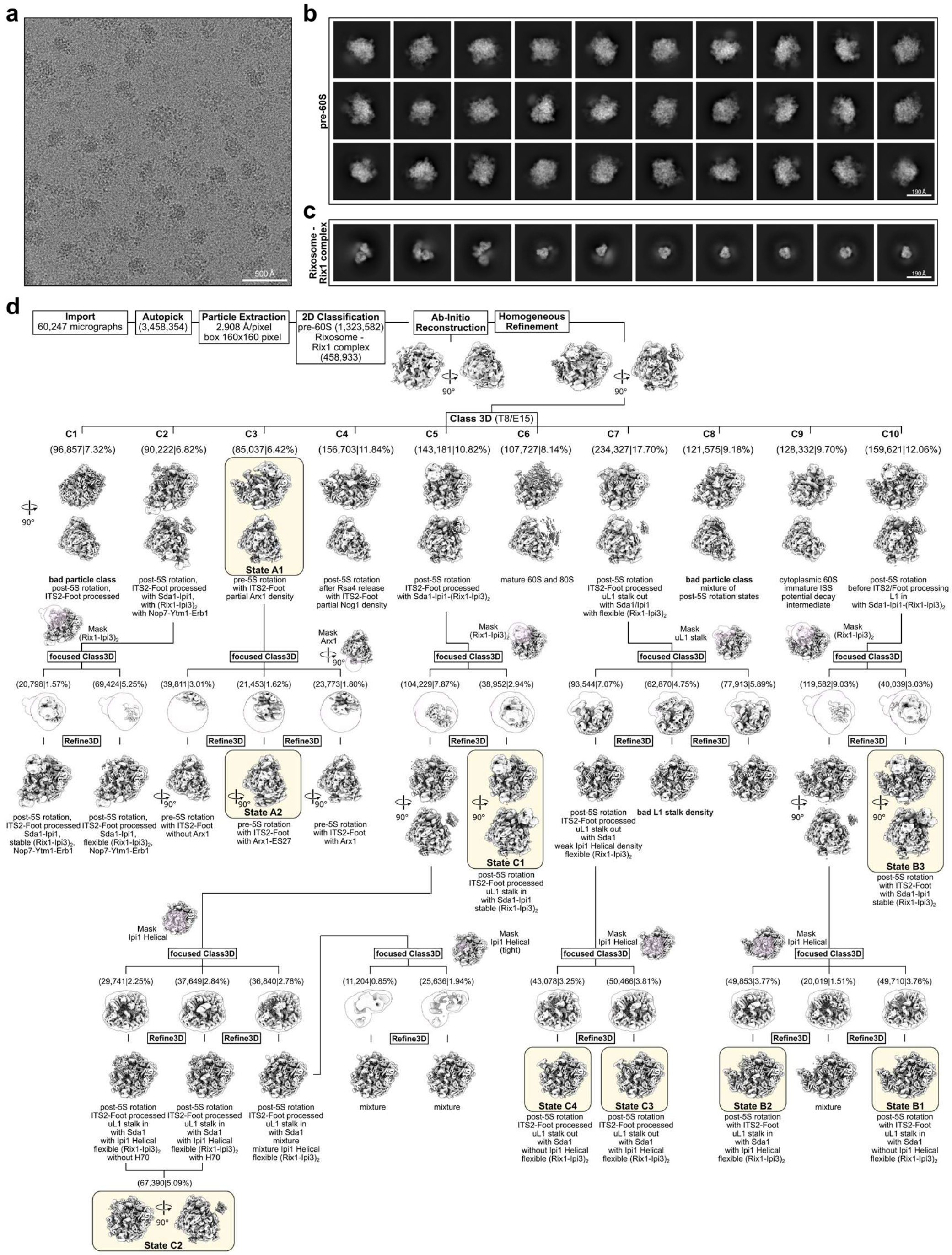

**Extended Data Fig. 11 | Cryo-EM analysis of Rix1-bound pre-60S ribosomes from *S.pombe*. a**, Representative cryo-EM micrograph of the Rix1 dataset. **b**-**c**, representative 2D class averages of Rix1-bound pre-60S subunits (**b**), and Rixosome and Rix1 complex particles (**c**). **d**, Cryo-EM sorting workflow for the classification of Rix1-bound pre-60S ribosomes. Extensive 3D classification resolved multiple pre-60S states, which were further refined and grouped into states A, B and C according to their maturation features. Final states are highlighted and particle numbers are shown in brackets.

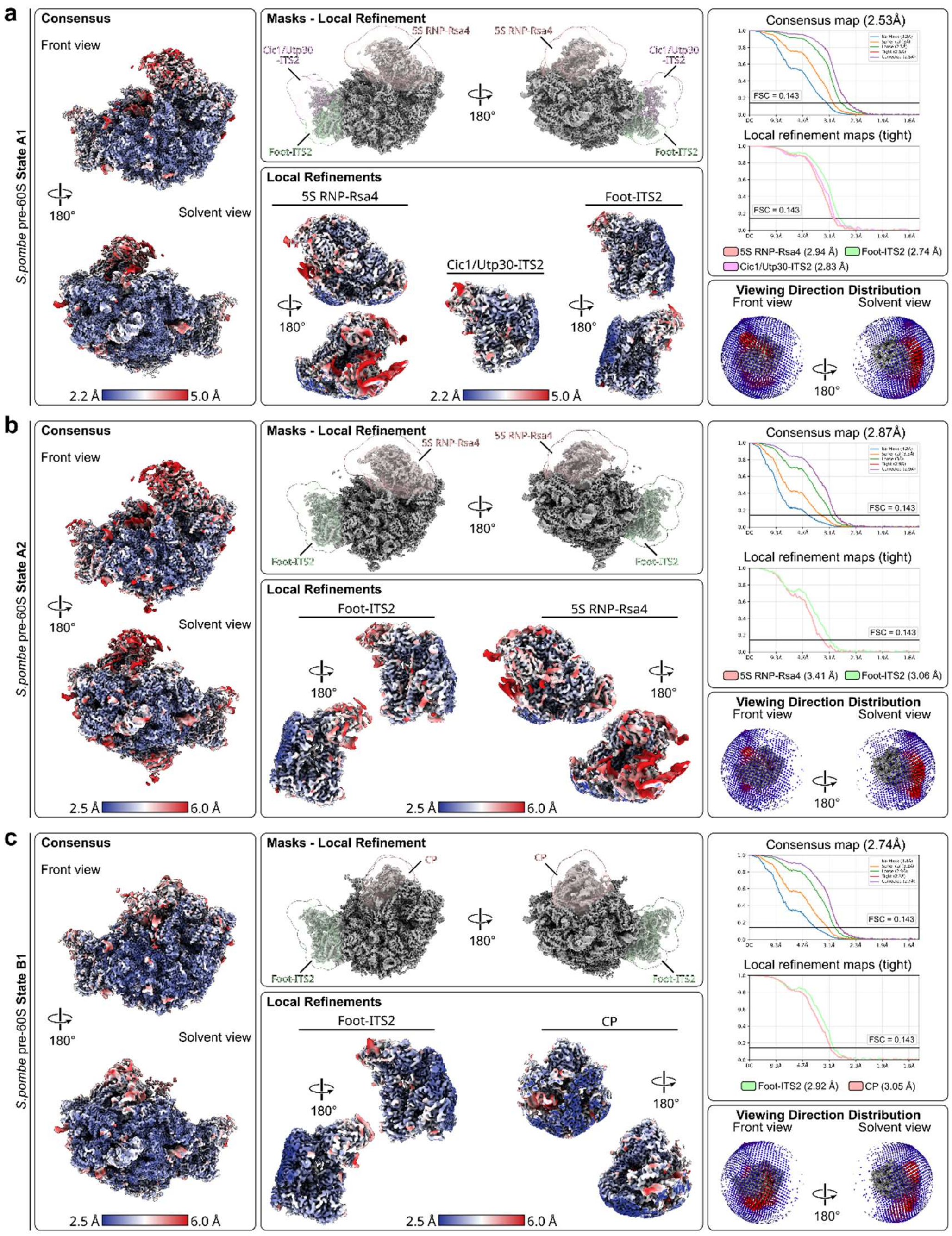

**Extended Data Fig. 12 | Cryo-EM validation of pre-60S ribosomal subunits from *S.pombe* (part 1). a-c**, Cryo-EM validation of the pre-60S state A1 (**a**), state A2 (**b**) and state B1 (**c**) reconstructions, including local resolution colored maps, masks used for local refinements, local refinement maps, Fourier shell correlation (FSC) curves and orientation distributions are shown.

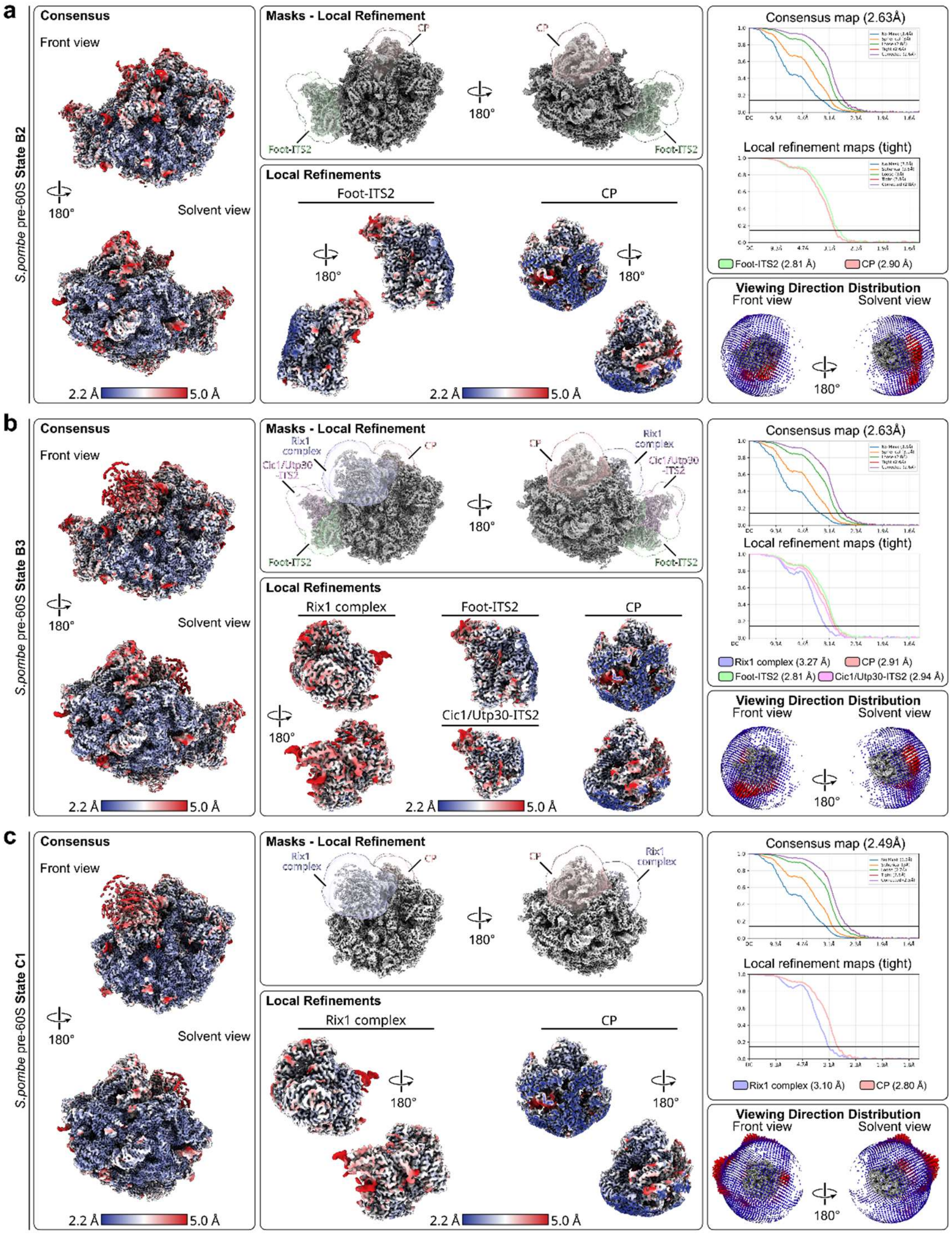

**Extended Data Fig. 13 | Cryo-EM validation of pre-60S ribosomal subunits from *S.pombe* (part 2). a-c**, Cryo-EM validation of the pre-60S state B2 (**a**), state B3 (**b**) and state C1 (**c**) reconstructions, including local resolution maps, masks used for local refinements, local refinement maps, Fourier shell correlation (FSC) curves and orientation distributions are shown.

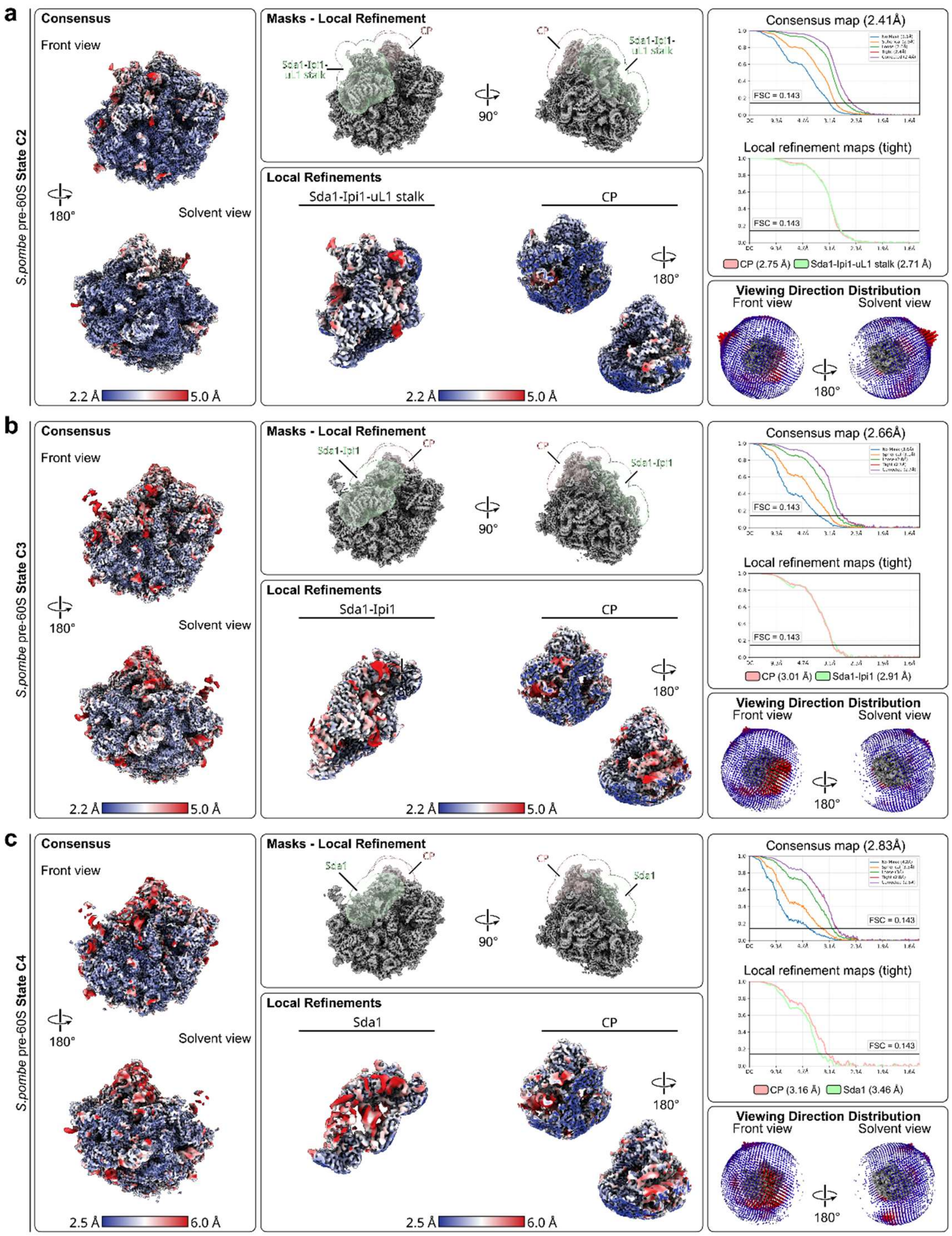

**Extended Data Fig. 14 | Cryo-EM validation of pre-60S ribosomal subunits from *S.pombe* (part 3). a-c**, Cryo-EM validation of the pre-60S state C2 (**a**), state C3 (**b**) and state C4 (**c**) reconstructions, including local resolution maps, masks used for local refinements, local refinement maps, Fourier shell correlation (FSC) curves and orientation distributions are shown.

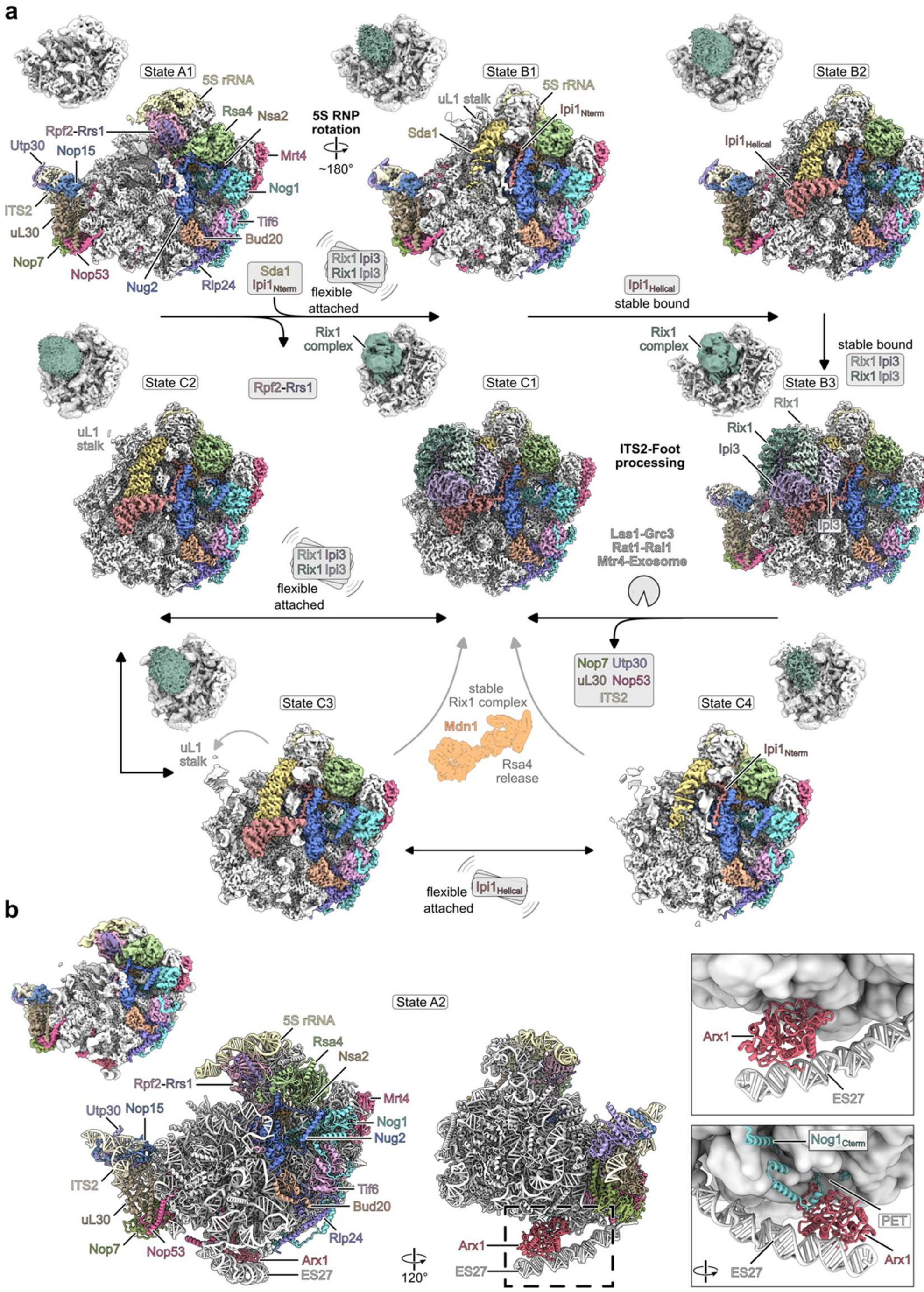

**Extended Data Fig. 15 | Cryo-EM analysis of pre-60S states from *S.pombe*. a,** Cryo-EM maps of the *S.pombe* pre-60S states purified through Rix1-FTpA. Assembly factors are colored individually. Low-pass filtered maps are shown in the upper corner of each state to visualize the Rix1 complex density (dark green). Arrows indicate the proposed maturation trajectory during ITS2 processing. Double headed arrows indicate that corresponding states may interconvert, whereas gray arrows indicate possible alternative maturation pathways. **b**, Cryo-EM map and atomic model of state A2. The close-up views show Arx1 bound to the polypeptide exit tunnel (PET) and in contact with expansion segment 27 (ES27) of the 18S rRNA and Nog1.

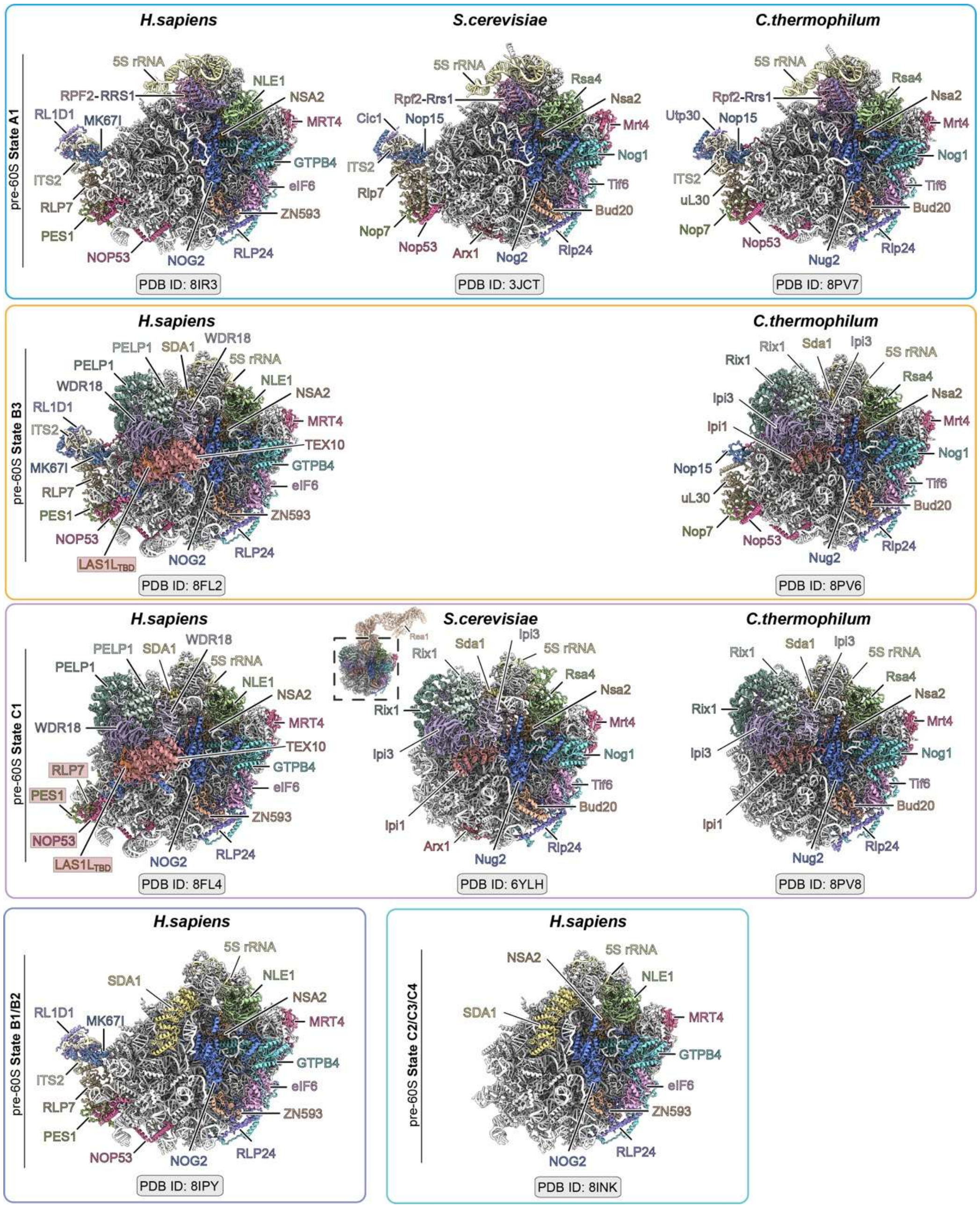

**Extended Data Fig. 16 | Comparison of pre-60S structures across species.** Similar pre-60S structures from *H. sapiens*, *S. cerevisiae* and *C. thermophilum* are shown for comparison with the *S. pombe* states described in this study. The structures are grouped according to related maturation stages, including early 5S RNP pre-rotate state corresponding to state A1, Rix1/RIX1 associated states corresponding to state B3, foot-removed states corresponding to state C1, and later human pre-60S states related to states B1/B2 or C2/C3/C4. Major assembly factors, ribosomal proteins and rRNA elements are labeled. PDB accession codes are indicated below each structure.

**Extended Data Table 1 | Cryo-EM data collection, refinement and validation statistics of human LAS1L-NOL9 dataset**

| | Human LAS1L-NOL9 |
|---|---|
| **Data collection and processing** | |
| Magnification | 130,000 |
| Voltage (kV) | 300 |
| Electron exposure (e–/Å$^2$) | ~50 |
| Defocus range (μm) | -1 to -2.5 |
| Pixel size (Å) | 0.932 |
| Symmetry imposed | *C1* |
| Initial particle images (no.) | 2,261,561 |
| Final particle images (no.) | 84043 |
| Map resolution (Å) | 3.4 |
| FSC threshold | 0.143 |
| Map resolution range (Å) | 2.9-10 |
| **Refinement** | |
| Model resolution (Å) | 3.5 |
| FSC threshold | 0.5 |
| Map sharpening *B* factor (Å$^2$) | -153 |
| Model composition | |
| Non-hydrogen atoms | 11504 |
| Protein residues | 1444 |
| *B* factors (Å$^2$) | 87.34 |
| Protein | 87.34 |
| R.m.s. deviations | |
| Bond lengths (Å) | 0.006 |
| Bond angles (°) | 1.166 |
| Validation | |
| MolProbity score | 2.00 |
| Clash score | 8.45 |
| Poor rotamers (%) | 1.69 |
| Ramachandran plot | |
| Favored (%) | 94.79 |
| Allowed (%) | 5.07 |
| Disallowed (%) | 0.14 |
| EMDB | XXXXX |
| PDB | XXXX |

**Extended Data Table 2 | Cryo-EM data collection, refinement and validation statistics of the Grc3 dataset**

| | *S.pombe* Rixosome (single Rix1 complex) | *S.pombe* Rixosome (double Rix1 complex) |
|---|---|---|
| **Data collection and processing** | | |
| Magnification | 165,000 | 165,000 |
| Voltage (kV) | 300 | 300 |
| Electron exposure (e–/Å$^2$) | 40 | 40 |
| Defocus range (μm) | 0.5-3.5 | 0.5-3.5 |
| Pixel size (Å) | 0.727 | 0.727 |
| Symmetry imposed | *C1* | *C1* |
| Initial particle images (no.) | 3,614,528 | 3,614,528 |
| Final particle images (no.) | 147,461 | 148,509 |
| Map resolution (Å) | 2.76 | 2.94 |
| FSC threshold | 0.143 | 0.143 |
| Masked map resolution range (Å) | 1.56-41.91 | 1.55-45.94 |
| **Refinement** | | |
| Model resolution (Å) | 2.7 | 2.8 |
| FSC threshold | 0.5 | 0.5 |
| Map sharpening *B* factor (Å$^2$) | -82 | -85 |
| Model composition | | |
| Non-hydrogen atoms | 27,168 | 43,341 |
| Protein residues | 3,582 | 5,785 |
| *B* factors (Å$^2$) mean | | |
| Protein | 73.94 | 69.91 |
| R.m.s. deviations | | |
| Bond lengths (Å) | 0.04 | 0.04 |
| Bond angles (°) | 0.769 | 0.826 |
| Validation | | |
| MolProbity score | 1.33 | 1.35 |
| Clash score | 3.52 | 3.99 |
| Poor rotamers (%) | 1.37 | 1.22 |
| Ramachandran plot | | |
| Favored (%) | 97.61 | 97.51 |
| Allowed (%) | 2.30 | 2.44 |
| Disallowed (%) | 0.09 | 0.05 |
| Map vs. Model CC (mask) | 0.86 | 0.84 |
| EMDB | EMD-XXXXX | EMD- XXXXX |
| Consensus & Local Refinements | EMD- XXXXX | EMD- XXXXX |
| | EMD- XXXXX | EMD- XXXXX |
| EMDB | EMD- XXXXX | EMD- XXXXX |
| PDB | XXXX | XXXX |

**Extended Data Table 3 | Cryo-EM data collection, refinement and validation statistics of the Rix1 dataset**

| | *S.pombe* pre-60S – State A1 | *S.pombe* pre-60S – State A2 | *S.pombe* pre-60S – State B1 | *S.pombe* pre-60S – State B2 | *S.pombe* pre-60S – State B3 |
|---|---|---|---|---|---|
| **Data collection and processing** | | | | | |
| Magnification | 165,000 | 165,000 | 165,000 | 165,000 | 165,000 |
| Voltage (kV) | 300 | 300 | 300 | 300 | 300 |
| Electron exposure (e–/Å$^2$) | 40 | 40 | 40 | 40 | 40 |
| Defocus range (μm) | 0.5-3.5 | 0.5-3.5 | 0.5-3.5 | 0.5-3.5 | 0.5-3.5 |
| Pixel size (Å) | 0.727 | 0.727 | 0.727 | 0.727 | 0.727 |
| Symmetry imposed | *C1* | *C1* | *C1* | *C1* | *C1* |
| Initial particle images (no.) | 3,458,354 | 3,458,354 | 3,458,354 | 3,458,354 | 3,458,354 |
| Final particle images (no.) | 85,037 | 21,453 | 49,710 | 49,853 | 40,039 |
| Map resolution (Å) | 2.53 | 2.87 | 2.74 | 2.63 | 2.63 |
| FSC threshold | 0.143 | 0.143 | 0.143 | 0.143 | 0.143 |
| Masked map resolution range (Å) | 1.57-38.63 | 1.55-46.45 | 1.56-42.56 | 1.56-42.56 | 1.56-42.59 |
| **Refinement** | | | | | |
| Model resolution (Å) | 2.5 | 2.8 | 2.7 | 2.6 | 2.6 |
| FSC threshold | 0.5 | 0.5 | 0.5 | 0.5 | 0.5 |
| Map sharpening *B* factor (Å$^2$) | -57 | -45 | -55 | -55 | -51 |
| Model composition | | | | | |
| Non-hydrogen atoms | 157,707 | 161,276 | 163,153 | 165,605 | 180,378 |
| Protein residues | 10,628 | 10,627 | 11,013 | 11,319 | 13,170 |
| Nucleotide residues | 3,460 | 3,628 | 3,561 | 3,561 | 3,577 |
| Water | 2 | 2 | 2 | 2 | 2 |
| Ligands | 7 | 7 | 8 | 8 | 8 |
| B factors (Å$^2$) mean | | | | | |
| Protein | 58.43 | 82.56 | 74.53 | 64.26 | 67.80 |
| Nucleotide | 30.40 | 64.21 | 48.72 | 40.27 | 40.33 |
| Ligand | 17.17 | 31.48 | 43.71 | 38.46 | 37.32 |
| R.m.s. deviations | | | | | |
| Bond lengths (Å) | 0.003 | 0.004 | 0.005 | 0.006 | 0.005 |
| Bond angles (°) | 0.752 | 0.787 | 0.822 | 0.850 | 0.749 |
| Validation | | | | | |
| MolProbity score | 1.17 | 1.62 | 1.63 | 1.52 | 1.28 |
| Clash score | 3.40 | 3.88 | 3.61 | 3.37 | 2.76 |
| Poor rotamers (%) | 0.99 | 2.66 | 2.72 | 2.26 | 1.69 |
| Ramachandran plot | | | | | |
| Favored (%) | 97.82 | 97.39 | 97.22 | 97.37 | 97.82 |
| Allowed (%) | 2.16 | 2.59 | 2.77 | 2.61 | 2.14 |
| Disallowed (%) | 0.02 | 0.02 | 0.01 | 0.02 | 0.05 |
| Map vs. Model CC (mask) | 0.88 | 0.89 | 0.90 | 0.90 | 0.90 |
| EMDB | | | | | |
| Consensus & Local Refinements | EMD-XXX<br>EMD-XXX<br>EMD-XXX<br>EMD-XXX | EMD-XXX<br>EMD-XXX<br>EMD-XXX | EMD-XXX<br>EMD-XXX<br>EMD-XXX | EMD-XXX<br>EMD-XXX<br>EMD-XXX | EMD-XXX<br>EMD-XXX<br>EMD-XXX<br>EMD-XXX<br>EMD-XXX |
| EMDB | EMD-XXX | EMD-XXX | EMD-XXX | EMD-XXX | EMD-XXX |
| PDB | XXXX | XXXX | XXXX | XXXX | XXXX |

| | *S.pombe* pre-60S – State C1 | *S.pombe* pre-60S – State C2 | *S.pombe* pre-60S – State C3 | *S.pombe* pre-60S – State C4 |
|---|---|---|---|---|
| **Data collection and processing** | | | | |
| Magnification | 165,000 | 165,000 | 165,000 | 165,000 |
| Voltage (kV) | 300 | 300 | 300 | 300 |
| Electron exposure (e–/Å$^2$) | 40 | 40 | 40 | 40 |
| Defocus range (μm) | 0.5-3.5 | 0.5-3.5 | 0.5-3.5 | 0.5-3.5 |
| Pixel size (Å) | 0.727 | 0.727 | 0.727 | 0.727 |
| Symmetry imposed | *C1* | *C1* | *C1* | *C1* |
| Initial particle images (no.) | 3,458,354 | 3,458,354 | 3,458,354 | 3,458,354 |
| Final particle images (no.) | 38,952 | 67,389 | 50,466 | 43,078 |
| Map resolution (Å) | 2.49 | 2.41 | 2.66 | 2.83 |
| FSC threshold | 0.143 | 0.143 | 0.143 | 0.143 |
| Masked map resolution range (Å) | 1.57-38.71 | 1.57-38.67 | 1.56-42.59 | 1.55-46.42 |
| **Refinement** | | | | |
| Model resolution (Å) | 2.6 | 2.3 | 2.8 | 2.8 |
| FSC threshold | 0.5 | 0.5 | 0.5 | 0.5 |
| Map sharpening *B* factor (Å$^2$) | -53 | -55 | -56 | -55 |
| Model composition | | | | |
| Non-hydrogen atoms | 167,215 | 153,591 | 153,591 | 150,061 |
| Protein residues | 11,851 | 10,000 | 10,000 | 9,671 |
| Nucleotide residues | 3,459 | 3,498 | 3,498 | 3,454 |
| Water | 2 | 2 | 2 | 2 |
| Ligands | 8 | 8 | 8 | 8 |
| B factors (Å$^2$) mean | | | | |
| Protein | 57.07 | 47.56 | 58.31 | 71.64 |
| Nucleotide | 34.51 | 32.56 | 38.74 | 31.84 |
| Ligand | 27.69 | 29.45 | 39.42 | 45.53 |
| R.m.s. deviations | | | | |
| Bond lengths (Å) | 0.010 | 0.004 | 0.004 | 0.004 |
| Bond angles (°) | 0.953 | 0.765 | 0.755 | 0.822 |
| Validation | | | | |
| MolProbity score | 1.47 | 1.49 | 1.53 | 1.61 |
| Clash score | 2.92 | 3.30 | 3.53 | 4.14 |
| Poor rotamers (%) | 1.66 | 2.12 | 2.06 | 2.34 |
| Ramachandran plot | | | | |
| Favored (%) | 96.54 | 97.40 | 97.27 | 97.33 |
| Allowed (%) | 3.40 | 2.55 | 2.78 | 2.65 |
| Disallowed (%) | 0.06 | 0.05 | 0.04 | 0.02 |
| Map vs. Model CC (mask) | 0.91 | 0.89 | 0.88 | 0.86 |
| EMDB | EMD-XXX | EMD-XXX | EMD-XXX | EMD-XXX |
| Consensus & Local Refinements | EMD-XXX | EMD-XXX | EMD-XXX | EMD-XXX |
| | EMD-XXX | EMD-XXX | EMD-XXX | EMD-XXX |
| EMDB | EMD-XXX | EMD-XXX | EMD-XXX | EMD-XXX |
| PDB | XXXX | XXXX | XXXX | XXXX |
